# Rat somatic genome editing enables ER+ breast cancer modeling

**DOI:** 10.1101/2025.09.15.675961

**Authors:** Wen Bu, Tobie Lee, Meenakshi Anurag, Yunfeng Ding, Ruixin Xu, Lillian He, Alexandria Z. Bu, Chandandeep Nagi, Carolina Gutierrez, Susan Hilsenbeck, Chonghui Cheng, Bing Zhang, Shixia Huang, Jianming Xu, C. Kent Osborne, Arun Sreekumar, Eric Chang, Chad J. Creighton, Xiang Zhang, Yi Li

**Affiliations:** Lester and Sue Smith Breast Center, Baylor College of Medicine, Houston, TX, USA; Department of Medicine, Baylor College of Medicine, Houston, TX, USA; Department of Pathology, Baylor College of Medicine, Houston, TX, USA; Dan L Duncan Comprehensive Cancer Center, Baylor College of Medicine, Houston, TX, USA; Department of Molecular and Human Genetics, Baylor College of Medicine, Houston, TX, USA; Department of Molecular and Cellular Biology, Baylor College of Medicine, Houston, TX, USA

## Abstract

Genetically engineered mouse models have advanced cancer research but often fail to capture key features of certain human tumors. Rats, with distinct physiology and tumor biology, offer a powerful alternative, yet their use has been constrained by technical barriers to genome editing. Here, we report efficient somatic genome editing in rats, enabling both Indel and substitution mutations. We then apply this approach to model estrogen receptor (ER)-positive breast cancer, which accounts for ∼70% of human cases but remains poorly represented in mice. The resulting rat tumors reproduce hallmarks of human ER+ breast cancer, including ductal histology, hormone responsiveness, and immune-microenvironmental features. By contrast, identical genetic alterations in mice failed to yield ER+ tumors, underscoring critical species differences in tumorigenesis. Together, this work establishes a versatile platform for rapid generation of clinically relevant rat tumor models, opening new avenues to dissect tumor biology, therapeutic response, and immune interactions in previously inaccessible cancer subtypes.

## Introduction

Rats, the first mammals domesticated for research, were instrumental in early cancer discoveries, including Charles Huggins’ demonstration of hormone dependency in prostate and breast cancer ^1^. However, enthusiasm for rats waned in the 1990s as genetic engineering advanced rapidly in mice, while similar techniques lagged in rats ^1,2^. Although mouse cancer models proliferated, they failed to mimic many human diseases. For example, *APC* mutation carriers develop colorectal cancer, but *Apc*-mutant mice develop tumors in the small intestine, whereas *Apc*-mutant rats develop colorectal tumors ^3^.

In breast cancer, hundreds of mouse models have advanced our understanding, yet they rarely represent estrogen receptor-positive (ER+) disease, the most common subtype, comprising approximately 70% of cases ^1,4^. ER+ tumors differ from HER2+ and triple-negative (lacking ER, progesterone receptor [PR, encoded by a transcriptional target of ER], and HER2) subtypes in their slower growth, well-differentiated histology, late bone-predominant recurrence, and responsiveness to endocrine therapy. Despite the high prevalence of ER+ cancers in humans, most mouse models lack ER and PR, and none require estrogen signaling for growth, except possibly Stat1 knockout mice ^1,2^. Consequently, ER+ tumors have been studied mainly in cell lines and xenografts, which lack intact immune microenvironments, critical in cancer development, progression and therapy.

Rats offer distinct advantages: their mammary gland architecture more closely resemble that of humans than mice, and carcinogen exposure readily induces ER+/PR+ tumors responsive to endocrine therapy ^1^. However, rat cancer models with defined genetic alterations remain scarce ^2^. The advent of CRISPR-Cas9 now enables efficient introduction of clinically relevant somatic mutations, enabling faithful modeling of sporadic cancers ^5^. While somatic genome editing has been applied to model breast cancer in mice ^6^, its potential in rats has yet to be unexplored.

Here, we adapt CRISPR-based somatic editing to rats, generating ER+ breast cancer models by mammary intraductal injection of adeno-associated virus (AAV) into Cas9-transgenic rats. AAVs carrying gRNAs (+ templates for homologous directed repair [HDR] as needed) introduced oncogenic alterations common in human ER+ breast cancer, producing tumors that express ER and PR, depend on estrogen signaling, and respond to endocrine therapy. Strikingly, identical editing in mice yielded ER-tumors, underscoring fundamental interspecies differences. These new rat models provide a long-missing system to study ER+ breast cancer within an intact immune context and establish a versatile platform for modeling cancers inadequately represented in mice.

## Results

### Intraductal AAV-gRNA delivery enables somatic genome editing in rats

We have reported efficient somatic genome editing in mouse mammary gland, generating several new mammary tumor models ^6^. Specifically, we intraductally injected adeno-associated virus (AAV)2/9 – AAV-2 inverted terminal repeat (ITR) and AA-9 capsid – to deliver gRNAs (and accessory components) into luminal epithelial cells in fully developed mammary glands of adult mice that are transgenic for Cas9^6^. Using this virus to edit breast cancer genes such as *Kras* and *Pik3ca*, we generated mammary tumors with high efficiency. To adapt this approach to rats, we first generated a rat line constitutively expressing *cas9* (Cas9 rat) by breeding LSL-Cas9 knock-in rats ^7^ with Cre transgenic rats ^8^. To test tropism of AAV serotype 9 in rat mammary gland epithelial cells, we intraductally delivered AAV2/9 virus carrying GFP into mammary glands of female Cas9 rats (8∼9 weeks; Figure S1A). At 3 days post injection, GFP was detected specifically in the infected glands based on fluorescent stereomicroscope (Figure S1B), immunohistochemical staining for GFP (IHC) (Figure S1C), and flow cytometry (Figure S1D&E).

To test whether this AAV vector can efficiently mediate genome editing in rats, we intraductally injected AAV virus (AAV-T) carrying a single guide RNA targeting exon 1 of *Tp53,* the most commonly mutated gene in human cancer. Of note, a higher amount of this virus (∼5x10^11^ gc) compared to AAV-GFP (∼2.5 × 10^11^ gc) was injected here to ensure adequate genome editing. Infected mammary glands were collected 4 days later, and the genomic DNA from the whole glands was extracted for PCR amplification of the target region. Next generation sequencing of the amplicon revealed Indel editing at approximately 4.2% of the sequenced alleles (Figure S2A & S2B), with the overwhelming majority (95%) being frameshift ^Indel^s (Figure S2C). Since the luminal epithelial cell proportion in these mammary glands is approximately 25∼30% of the total cell population ^9^, the genome editing rate of the luminal cell population is estimated to be 10.5-21% depending on whether editing occurred in one or both alleles in targeted cells. Together, these data demonstrate that our AAV-gRNA vector can efficiently edit somatic genome in rats.

### Somatic Indel editing in rats efficiently inactivates tumor suppressors and induces mammary tumorigenesis

Having confirmed efficient somatic genome editing, we next tested the efficacy of this virus in inducing mammary tumors. We injected AAV-T into ten 8∼9-week-old Cas9 females (∼5x10^11^gc in a single #4 gland per rat). Palpable tumors started to appear in approximately 3 months, and by 10 months, 50% of rats developed palpable tumors (Figure 1A & 1B). To confirm Indel editing in these tumors, we extracted tumor genomic DNA, PCR-amplified the target region and then performed amplicon deep sequencing.

**Figure 1.**
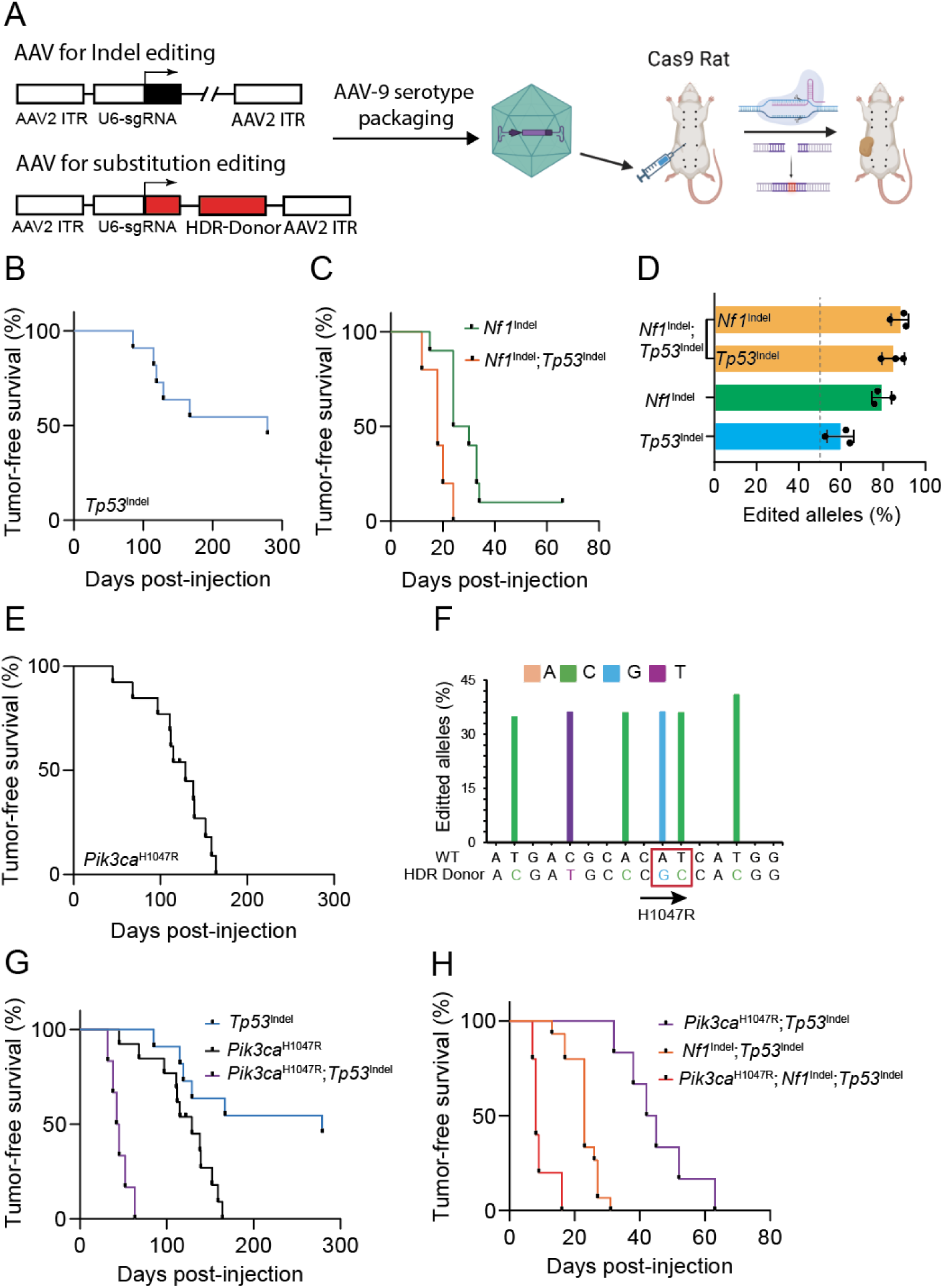
Intraductal injection of AAV vectors to deliver gRNA to edit genes in mammary glands to initiate tumorigenesis. A) Schematic representation of mammary tumor model generation via intraductal injection of AAV to edit mammary cells in rats. (B, C, E, G, and H) Kaplan-Meier tumor-free survival curves of rats injected intraductally with AAV to introduce *Tp53* indels (4.7x10^11^gc/gland, age=9∼13, n=10) (B)*, Nf1* indels (2.8x10^11^gc/gland, Age=10∼14 weeks, n=10) (C), *Tp53* indels + *Nf1* indels (2x10^10^gc/gland, age=10∼11 weeks, n=10) (C), *Pik3ca*^H1047R^ (4x10^11^gc/gland, age=9∼14, n=12) (E), Tp53 indels + *Pik3ca*^H1047R^ (5.9x10^10^gc/gland, age=11∼12weeks, n=6) (G), and *Tp53* indels + *Nf1* indels + *Pik3ca*^H1047R^ (1.43x10^10^gc/gland, age=10weeks, n=5) (H). Tumor latencies of rats with individual and double editing are shown for comparison in (G) and (H), respectively. (D and F) Bar graphs of percentages of edited alleles in tumors from (B), (C), and (E). In addition to the H1047R mutation in *Pik3ca*, four silent mutations were introduced to prevent re-editing of the HDR donor sequence.

Indel editing occurred at the expected sites (Figure S3A) and in approximately 60% of the sequenced genomic alleles (Figure S3A & 1C). Considering notable involvement of stroma (Fig 2) that contributed to the non-edited genomic copies, genome editing likely occurred in both alleles of at least most, if not all, of the mammary tumor cells. Biallelic deactivation is also likely needed for these mammary cells to evolve into cancer based on previous studies of *Tp53* in mice and in human breast cancer specimens ^10^. Of note, while editing may have occurred in only one allele among many of the initially infected cells, only biallelically mutated cells likely have competitive advantage in tumor evolution. Together, these data demonstrate successful tumor induction by somatic Indel editing of a tumor suppressor gene. Interestingly, the tumor latency of 10 months at median in these rats is comparable to that in mice similarly mutated for *Tp53* ^11^.

**Figure 2.**
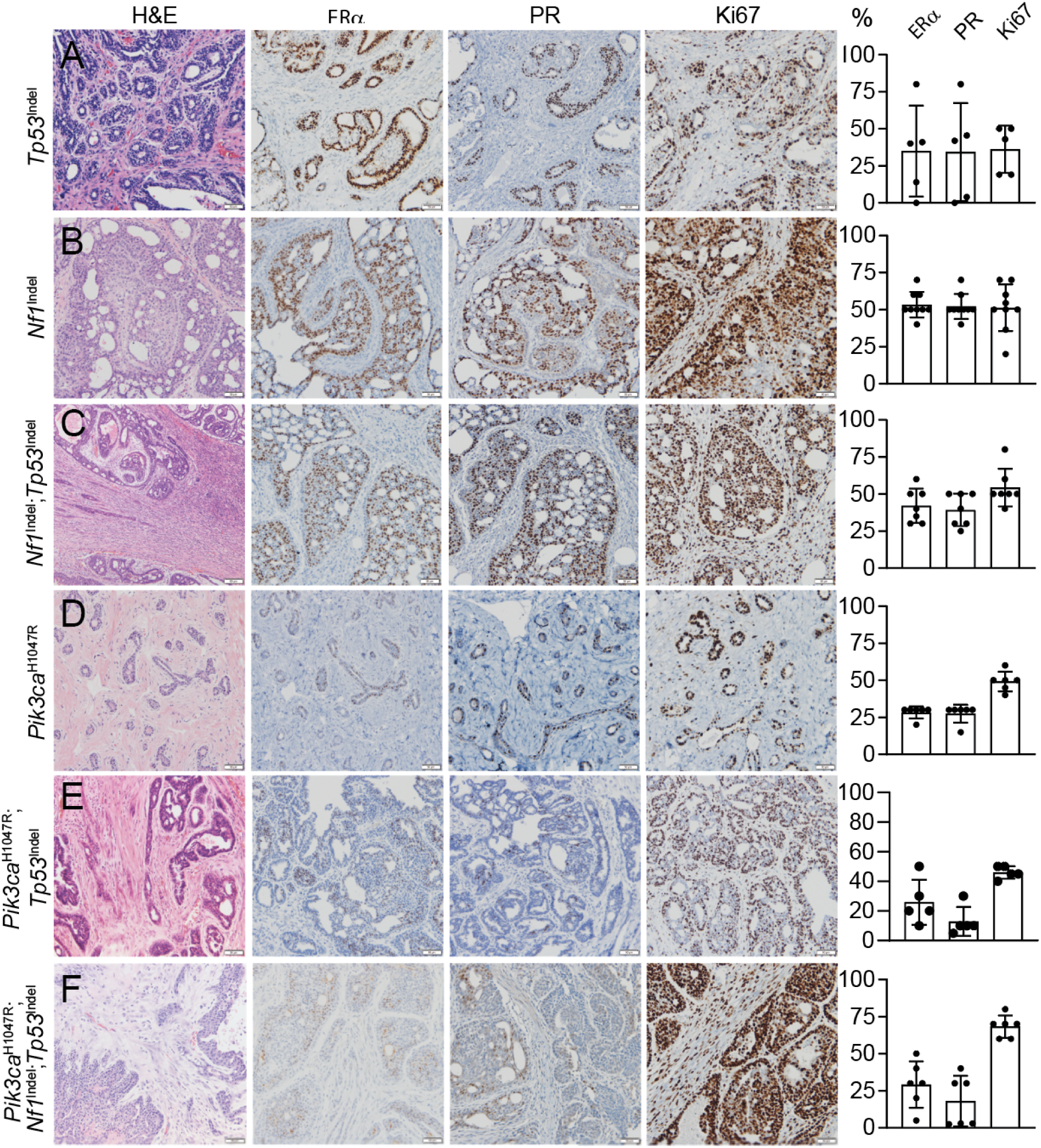
Histological and biomarker characterizations of mammary tumors induced by somatic genome editing. Representative photomicrographs of HE and IHC of mammary tumors induced by intraductal injection of AAV carrying gRNAs to introduce the indicated genetic alterations. The biomarkers detected by IHC are labeled at the top. The proportions of ER, PR and Ki67-postive tumor cells are shown in the bar graphs to the right of each row.

To further confirm this somatic editing approach for generating rat models of breast cancer, we targeted another tumor suppressor that is known to be involved in human ER+ breast cancer. *Nf1* encodes neurofibromin, a GAP (GTPase Activating Protein) that inactivates RAS and also acts as a co-repressor of ER-mediated gene transcription ^12^. In early-stage ER+ breast cancers, the frequencies of *Nf1* mutations and copy number loss are 4% and 20%, respectively ^12–14^. Intraductal injection into a mammary gland of AAV2/9 virus carrying a gRNA targeting *Nf1* gene (AAV-N; 2.8x10^11^gc/gland) led to palpable tumors with median latency of 4 weeks (Fig 1C). Indel editing occurred at the expected sites (Figure S3B) and in approximately 80% of genomic alleles (Figure S3B & 1D). Considering substantial involvement of stromal cells that were not edited, this 80% overall editing suggests that most, if not all, tumor cells were edited in both alleles, further demonstrating efficient tumor induction by AAV-mediated somatic Indel editing. The tumor latency in these rats is substantially faster than tumor latency in rats with a germline deletion of one copy of *Nf1* (18 days vs. 70-90 days depending on the line)^15^, suggesting that somatic mutations of *Nf1* are potent in initiating cancer and that homozygous mutations are likely more transforming than heterozygous mutations which may require loss of heterozygosity to complete tumorigenesis.

Since human cancers are often caused by more than one oncogenic mutation, we next tested whether we can concurrently edit two tumor suppressors for tumor induction. Since *Nf1* and *Tp5*3 are frequently co-mutated in human breast cancer ^14,16^, we made an AAV2/9 virus carrying 2 sgRNAs targeting *Nf1* gene and *Tp53* gene (AAV-NT). Intraductal delivery of AAV-NT (2x10^10^gc/gland) into 10 Cas9 rats led to tumors within one month for all injected rats (Figure 1C). Amplicon sequencing revealed Indel editing of each gene at 85-90% of genomic copies of these tumors (Fig S3C & 1D), suggesting that both alleles of both genes are likely mutated in all tumor cells, demonstrating the high efficacy of our vector system to edit two genes concurrently to initiate tumorigenesis. Of note, the moderately higher percentage of edited genomic copies in these tumors compared to tumors initiated by *Tp53* alone is likely due to less stromal cell infiltration in these double mutated tumors. In addition, the significantly shorter tumor latency in these rats compared to that in rats injected with either AAV-N or AAV-T (p<0.01) demonstrates an oncogenic collaboration between these *Nf1* and *Tp53* mutations in driving breast tumorigenesis.

### Somatic substitution editing in rats effectively activates protooncogenes and induces tumorigenesis

Besides loss-of-function mutations of tumor suppressor genes, gain-of-function mutations of protooncogenes are the other key driver of tumorigenesis in breast and other organs. We have previously optimized an AAV vector for HDR-based precision-editing of protooncogenes in mice to initiate tumorigenesis ^6^. Especially, we demonstrated efficient tumorigenesis in mice intraductally injected with AAV to edit *Pik3ca* to *Pik3ca^H1047R^*. *PIK3CA* is the most commonly mutated protooncogene in human breast cancer, and H1047R is the most frequent mutation in *PIK3CA*, leading to constitutive activation of PI3K signaling ^17^. Therefore, in this vector, we cloned sgRNA and HDR donor sequence targeting rat *Pik3ca* for the H1047R mutation (Fig 1A). Intraductal injection of the resulting virus, AAV-P, led to mammary tumors with a median latency of 4.5 months (Figure 1E). Amplicon sequencing of the edited region detected the H1047R mutation in approximately 40% of the genomic alleles of the tumors. Similar frequencies of silent genomic changes were detected near H1047R (Figure 1F). These silent mutations were designed to prevent HDR from being targeted by sgRNA/Cas9. Together, these data demonstrate successful substitution editing by our AAV viral vector. The observation of ∼40% editing of the *Pik3ca* alleles, compared with ∼80% of the *Nf1* alleles in tumors induced by AAV-N, likely reflects the distinct functional requirements of these gene classes. Whereas oncogene activation typically requires mutation of only a single allele to drive tumorigenesis, effective inactivation of a tumor suppressor generally necessitates disruption of both alleles.

### Somatic editing of both tumor suppressors and protooncogenes in rats rapidly initiates tumorigenesis

Human tumors are often caused by a combination of mutated tumor suppressors and activated protooncogenes. To test whether our vector system could efficiently introduce both Indel and substitution editing in mammary cells in rats, we attempted to Indel-edit *Tp53* and substitution -edit *Pik3ca*, two genes that are known to be co-mutated in substantial proportions of human breast cancers (10.5%; METABRIC), and have been reported to collaborate in transforming human and mouse mammary cells ^18,19^ and cause worse prognosis ^20^. To this end, we moved the U6 promoter-driven sgRNA targeting *Tp53* into AAV-P vector. Intraductal injection of the resulting virus (AAV-PT) into Cas9 rats led to tumors with a median latency of only 42 days, markedly faster than targeting either *Tp53* or *Pik3ca* alone (Fig 1G), confirming the high synergy of these two most common breast cancer genes.

In addition, we tested whether our vector system could concurrently genome-edit multiple genes. This is important since human cancers are often caused by more than two genetic mutations. Therefore, we built an AAV vector that targets *TP53*, *NF1* and *PIK3CA.* Intraductal injection of AAV-PNT into Cas9 rats led to tumors with an exceedingly short median latency of only 8 days (Figure 1H), significantly faster than either *Pik3ca*^H1047R^/*Tp53*^Indel^ tumors or *Nf1*^Indel^ /*Tp53*^Indel^ tumors, demonstrating the collaboration of these three genetic lesions in breast tumorigenesis.

### Somatic genome editing-induced mammary tumors in rats are ER^+^PR^+^ and exhibit from benign to high grade histological features

Having generated a panel of rat models of breast cancer, we examined their histology and hormone receptor expression. In women, 75-80% of breast cancers are ductal carcinomas, while 10-15% are lobular carcinomas which usually carry E-cadherin mutations and need to be modeled in animals with E-cadherin loss. Among ductal carcinomas, most (75-80%) are not otherwise specified (NOS), with only 10-15% representing special histological subtypes such as mucinous, medullary, papillary or metaplastic variants. Approximately 70% of ductal carcinomas express ER, often along with PR. Histologically, our six groups of rat tumors displayed two notable features: robust hormone receptor expression and an absence of squamous differentiation. This closely mirrors the majority of human ductal carcinomas but diverges from most mouse models, which are overwhelmingly ER-/PR- and frequently exhibit squamous differentiation ^1^.

All five tumors induced by *Tp53*^Indel^-editing were ductal carcinomas (Fig. 2A); but one of them contained moderate areas of sarcomatous and osteosarcomatous differentiation, and another exhibited sarcoma/fibrosarcoma with focal liposarcoma features (Fig. S4). ER/PR expression ranged from absent to >75%, likely due to divergent secondary spontaneous mutations (Fig. 2A). In contrast, mouse *Tp53*^Indel^ tumors lack ER/PR and show squamous and spindle differentiation ^11^

Tumors induced by *Nf1* Indel-editing were predominantly ductal carcinoma in situ (12/13 cases; the remaining two case were a benign papilloma and an invasive ductal carcinoma), all with moderate differentiation and high similarity to one another. These tumors contained moderate to high proportions of ER+ cells and Ki67+ cells (Fig. 2B). Histologically, they resembled ER+/PR+ tumors reported in *Nf1* germline mutant rats ^15^, although those germline tumors were reported to be invasive.

By contrast, combined *Nf1* and *Tp53* mutations yielded invasive ductal carcinomas with moderate differentiation in all eight cases examined (Fig. 2C), suggesting that *Tp53* loss promotes invasion of *Nf1*^Indel^-initiated lesions. These findings also imply that *Nf1* inactivation restrains *Tp53*-mutated cells from undergoing metaplastic differentiation. Like *Nf1*^Indel^-only tumors, the double mutants contained moderate to high percentages of ER+ cells and Ki67+ cells (Fig. 2C).

Introduction of the *Pik3ca*^H1047R^ mutation produced benign fibroadenomas in 8 of 10 cases (the remainder being a lactational adenoma and a ductal carcinoma NOS) (Fig. 2D). *Pik3ca* mutations have also been identified in human breast fibroepithelial tumors ^21^. ER was detected in ∼27% of ductal cells, similar to the 10-30% reported in normal rat terminal duct lobular units ^22^. Ki67 was present in ∼50% of epithelial tumor cells and in the fibrous component, reflecting the high proliferative activity of both compartments.

In contrast, combined *Tp53*^Indel^ and *Pik3ca*^H1047R^ mutations produced moderately differentiated invasive ductal carcinomas with papillary features in all five cases (Fig. 2E). This suggests that *Tp53* loss drives progression from fibroadenoma to carcinoma, while PIK3CA activation restricts *Tp53*-mutated cells from transdifferentiation, unlike *Tp53*^Indel^-only tumors. These tumors contained moderate levels of ER+ cells but relatively low PR+ expression.

Finally, triple mutations (*Tp53*^Indel^ + *Nf1*^Indel^ + *Pik3ca*^H1047R^) produced invasive ductal carcinomas with moderate to poor differentiation, nuclear pleomorphism, high proliferation, and extensive necrosis in all seven cases (Fig. 2F). These tumors were of higher grade than either *Tp53*^Indel^ + *Pik3ca*^H1047R^ or *Tp53*^Indel^ + *Nf1*^Indel^ tumors, indicating the additive progression impact of a third oncogenic driver. ER expression was moderate, while PR expression was relatively low (Fig. 2F).

### Somatic genome editing-induced mammary tumor models in rats show overlapping and distinct transcriptome profiles

After having characterized the histology and basic biomarkers of these rat ER+/PR+ tumors, we next compared their transcriptomes. ANOVA analysis (FDR<1%) identified 1,579 differentially expressed genes across 35 rat tumors in our 6 models. The inferred expression-based intrinsic subtypes of most tumors were Luminal A or Luminal B, consistent with positive IHC for ER, PR, and Ki67. However, two each of *Tp53*^Indel^ tumors, *Pik3ca*^H1047R^/*Tp53*^Indel^ tumors, and *Pik3ca*^H1047R^/*Tp53*^Indel^/*Nf1*^Indel^ tumors were mapped to Basal-like although they produce detectable ER and PR protein. Two tumors (one in *Nf1*^Indel^/*Tp53*^Indel^ and one in *Pik3ca*^H1047R^/*Tp53*^Indel^) were mapped to HER2 subtype though these did not correspond to elevated HER2 mRNA (Fig 3A).

**Figure 3.**
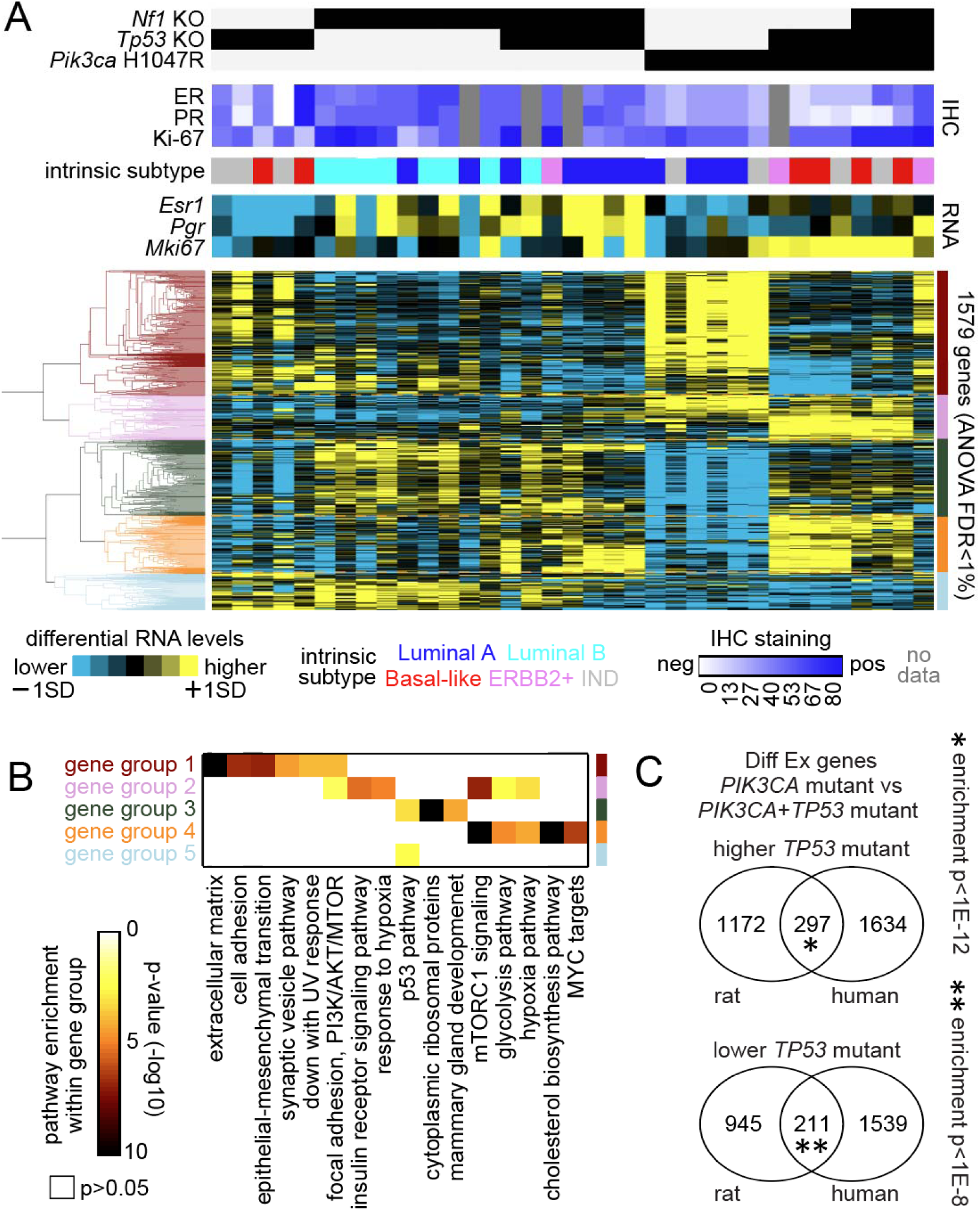
Molecular heterogeneity across rat tumor models according to genotype. (a) Across 34 rat model breast tumors, differential expression patterns for the top genes differing according to rat model genotype (FDR<1%, ANOVA), involving six genotypes in total. From unsupervised clustering of the genes, five major gene groups are evident. Rat tumors were assayed for ER and PR protein levels by IHC and also categorized into intrinsic expression-based subtypes using RNA-seq data (IND, indeterminate subtype call). (b) Significance of enrichment (by one-sided Fisher’s exact test) for selected Hallmark pathway or Gene Ontology gene sets within each of the five differential gene sets as defined in part a. (c) Genes differentially expressed in rat *Pik3ca^H1014R^*+*TP53* wt versus *Pik3ca^H1047R^*+*Tp53*^Indel^ tumors (using p<0.01, t-test using log2-transformed data) shared significant overlap with the gene orthologs arising from the analogous comparison in human breast tumors (p<0.001, t-test). Human breast tumor comparisons are based on 317 tumors harboring a hotspot mutation in PIK3CA. P-values by one-sided Fisher’s exact tests.

Hierarchical clustering using these 1,579 DEGs shows extensive heterogeneity across models, as expected by their diverse oncogenic drivers and histological features (Figure 3A and 3B). One group of genes (“group 1”)—enriched prominently for extracellular matrix, cell adhesion and EMT genes—was differentially higher in *Pik3ca*^H1047R^ tumors, consistent with the presence of large amounts of stroma tissues in these tumors. This gene group was also higher in two of the 5 *Tp53*^Indel^ tumors, highlighting the heterogeneous nature of these *Tp53*^Indel^ tumors. On the other hand, another group of genes (“group 2”), enriched for insulin receptor signaling and mTOR signaling genes, was higher in all three models that include *Pik3ca*^H1047R^ as an initiating oncogene, likely reflecting PI3K signaling in driving insulin and mTOR signaling. In contrast, despite *Nf1*^Indel^ alone being more potent tumor inducer than *Pik3ca*^H1047R^ by itself, no discernible gene group emerged across the models that harbor a *Nf1* loss of function mutation. Since *Tp53*^Indel^ is a poor driver of cancer on its own, not surprisingly, the three tumor models that harbored *Tp53*^Indel^ did not share a distinct gene group. In accordance with this finding, one gene group ("group 5”), modestly enriched for the p53 pathway, was higher in *Tp53*^Indel^ tumors and *Nf1*^Indel^/*Tp53*^Indel^ tumors but even higher in *Nf1*^Indel^ tumors and lower in two other models that harbored *Tp53*^Indel^, again revealing the non-dominant nature of *Tp53* mutations in defining gene expression profiles of the eventual tumors that have likely gained other and more dominant oncogenic drivers.

The gene “group 3”, enriched for cytoplasmic ribosomal genes and mammary gland development-related genes, was higher in *Nf1*^Indel^ tumors and *Nf1*^Indel^/*Tp53*^Indel^ tumors, especially those expressing high levels of *Esr1* and *Pgr1*, consistent with Ras on translational regulation and ER on mammary gland development. However, this gene group was not elevated in the *Pik3ca*^H1047R^/*Tp53*^Indel^/*Nf1*^Indel^ tumors, suggesting that *Pik3ca*^H1047R^ alone or together with *Tp53* mutations may mask some of the signal output from *Nf1* loss. Interestingly, this gene group was depleted among the *Pik3ca*^H1047^ and the *Tp53*^Indel^ tumors exhibiting robust “group 1” genes, likely due to the non-luminal cell component in this latter two models diluting high cytoplasmic ribosomal activities in the luminal cell component and driving genes involved in extracellular matrix, cell adhesion, and EMT.

The gene “group 4” —enriched for multiple pathways including mTOR signaling, cholesterol biosynthesis pathway, and MYC targets as well as to a lesser degree glycolysis pathway—was robust in *Pik3ca*^H1047R^/*Tp53*^Indel^ tumors, and to a lesser degree in the *Nf1*^Indel^/*Tp53*^Indel^ tumors and *Pik3ca*^H1047R^/*Tp53*^Indel^/*Nf1*^Indel^ tumors, suggestive of increased metabolism in these tumors. This gene group conceivably represents the output of the combinatorial effects of *Tp53* loss and PI3K activation since it did not manifest in other tumor models that harbored *Tp53*^Indel^ or *Pik3ca*^H1047^ alone. Although *Pik3ca* itself is not mutated in *Nf1*^Indel^/*Tp53*^Indel^ tumors, *Nf1* loss activates Ras which activates PI3K signaling. Together, these transcriptome data confirm the heterogeneous nature of these tumor models and reveal mutation-driven signaling pathways that can be heavily influenced/masked by co-mutations.

To determine how well these rat tumors mimic human breast cancer, we compared their transcriptome of the *Pik3ca*^H1047R^/*Tp53*^Indel^ model with the human breast cancers also bearing these two mutations. We chose this rat model for comparison because these two gene mutations are the two most common genetic alterations in human ER+ breast cancer as well as human breast cancer in general and because these two genes drive rapid formation of ductal carcinoma, the most common pathological type of human breast cancer. As shown in Fig 3C, among the top 1,469 genes higher in *Tp53*^Indel^/*Pik3ca*^H1047R^ tumors over *Pik3ca*^H1047R^ only tumors, 297 genes are also enriched in genes higher in TCGA breast cancer cases mutated for both *TP53* (loss or point mutations) and *PIK3CA* (point mutations) when compared to cases mutated for *PIK3CA* but WT for *TP53* (enrichment p value <1E-14, one-sided Fisher’s exact test).

Likewise, among 1,156 genes lower in the rat comparison, 211 genes are also lower in the TCGA comparison (enrichment p <1E-6). Together, these data demonstrate that these rat tumors significantly recapitulate gene expression changes in patients as a result of *Tp53*^Indel^ changes.

### Genome editing-induced mammary tumors in rats are distinct from tumors in mice

While all of these CRISPR rat models develop ER+ ductal tumors (or fibroadenoma in the *Pik3ca*^H1047R^ rats) with little metaplastic differentiation, very few of the existing many mouse models develop similar tumors. In fact, ER negativity and/or metaplastic changes, especially squamous differentiation, are a common feature of some of the most common mouse models of breast cancer induced by a variety of oncogenic drivers ^23^. While some of the genetic drivers in mouse models have also been introduced into rats, transgenic lines between species or even within the same species cannot be properly compared due to the randomness of transgene insertions into the genome and variations of transgene copy number inserted, leading to large differences in gene expression levels between rodent lines. Germline knockout lines between species can be more properly compared, but the few relevant rat lines such as *Tp53*, *Brca1*, and *Brca2* knockouts failed to develop mammary tumors on their own ^2^. Therefore, we considered somatic models. We have reported that intraductal injection of the lentivirus carrying mouse *Hras*^Q61L^ into mice led to ER-, metaplastic mammary tumors with heavy squamous differentiation ^1^, but the same virus led to ER+/PR+ ductal carcinomas in rats ^1,24^. However, because lentiviral overexpression relies on random insertion and ectopic expression, these models carry inherent limitations and leave unresolved concerns in comparing mouse versus rat models.

Somatic CRISPR genome editing minimizes potential pitfalls associated with other methods to deliver oncogenic lesions and offers a direct comparison between mice and rats. In our recent publication of somatic genomic editing in mice, we used intraductal injection of AAV to edit *Kras* to *Kras*^G12D^ and generated mammary tumors that are characterized as ER-/PR-ductal carcinoma with widespread and extensive squamous differentiation ^6^, histologically similar to the mouse tumors induced by intraductal injection of lenti-HrasQ61L ^6^. Therefore, we compared this mouse model to tumors in rats induced by an identical method. Intraductal injection into rats of AAV carrying gRNA to introduce G12D into *Kras* led to palpable tumors with a median short latency of approximately 70 days. However, these *Kras*^G12D^ rat tumors are all ER+/PR+ ductal carcinoma with papillary features but no metaplasia (Fig 4A), similar to the tumors induced by intraductal injection of lenti-*Hras*^Q61L^24. Therefore, regardless of whether Ras activation is achieved by CRISPR-mediated genome editing or lentivirus-mediated ectopic expression of an activation mutated, rats develop ER+/PR+ tumors, but mice form ER-/PR-metaplastic tumors with heavy squamous differentiation. These data highlight a dramatic difference between these two rodent species in mammary tumorigenesis.

**Figure 4.**
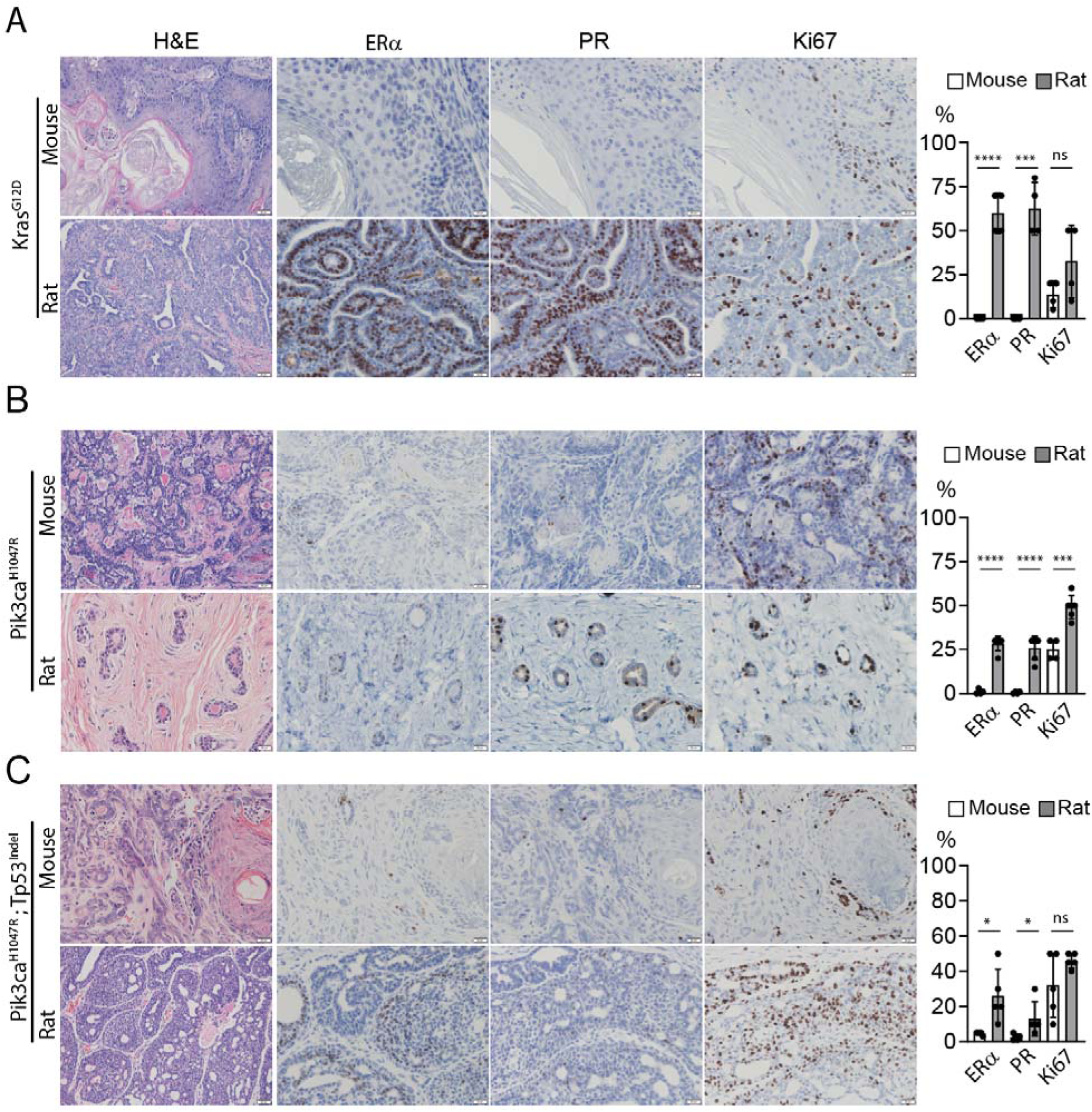
Comparisons of mouse and rat tumors induced by identical oncogenic drivers. Representative H&E and IHC staining of tumors induced by somatic genome editing of the indicated genes in the indicated host. The proportion of ER, PR, and Ki67 positive tumor cells are shown on the right.

*Kras* is not commonly mutated in human breast cancer, but *Pik3ca* is the most commonly mutated breast cancer proto-oncogenes. Therefore, we next compared *Pik3ca*^H1047R^ tumors in mice vs. rats. We have previously reported mouse tumors induced by intraductal injection of AAV to edit *Pik3ca* to *Pik3ca*^H1047R6^. The resulting tumors were detected with a latency of 4.9 months ^6^, similar to 4.3 months in rats injected by AAV-edited *Pik3ca* (Fig S5), but these mouse tumors are all characterized as low to moderate grade ductal carcinomas with focal squamous differentiation and negligible proportions of ER+ or PR+ cells (Fig 4B), dramatically different from those of rat lesions induced by genome editing of *Pik3ca*, which are benign fibroadenoma featuring normal-appearing ducts and without any metaplastic changes.

The relatively long latencies of these mouse and rat *Pik3ca*^H1047R^ tumors imply that they spontaneously gained secondary mutations, which might also have contributed to the final tumor histology. To minimize their involvement and complications, we next compared our rat tumors induced by the two most common and potent oncogenic drivers in ER+ breast cancer, *Pik3ca* and *Tp53* (Fig 2E) to the tumors in mice intraductally injected with AAV to edit the same two genes. The mouse tumors appeared with a short latency similar to that of the rat tumors (Figure S6). All five mouse tumors are poorly differentiated high-grade metaplastic carcinoma with predominant sarcomatous differentiation and focal squamous differentiation as well as with the residual adeno-appearing tumor cells occasionally expressing ER/PR (Fig 4C). These characteristics are drastically different from all five rat counterparts, which are moderately differentiated ER+/PR+ ductal carcinoma with papillary features, again revealing a striking difference in tumor characteristics between these two rodent species. Together, these data from three well-controlled comparisons between mice and rats demonstrate a dramatic divergence of these two rodent species in mammary tumor development—while mice predominantly develop mammary tumors with squamous metaplasia, a very rare subtype of human breast cancer, rats commonly develop ER+/PR+ ductal carcinoma NOS, the most common type of human breast cancer responsible for 70% of all human breast cancer cases, further demonstrating the value of these somatic rat models in providing a platform for studying the most common type of breast cancer in its native immune microenvironment.

### The *Nf1* Indel mammary tumor model is endocrine therapy-sensitive, while the *Nf1*/*Tp53* Indel mammary tumor model is endocrine therapy-resistant

Given that these rat tumors are ER+/PR+ and can be designated as Luminal A or Luminal B, we next tested their hormone dependency and therefore potential as preclinical models to test new endocrine therapies. We started with the *Nf1*^Indel^ model for its high levels of ER+/PR+ and for known involvement of *Nf1* loss in derepressing ER transcription and promoting endocrine therapy resistance in patients ^12^. Ovariectomy—which eliminates the primary source of estrogen although it spares the residual estrogen from extra-ovarian sources such as adipose tissue and adrenal glands—is used in certain high-risk patients and also as a surgical alternative to estrogen deprivation treatment such as aromatase inhibitors (AI) + ovarian suppression. Ovariectomy of 3 rats bearing *Nf1*^Indel^-induced tumors caused rapid and complete regression in all three animals, demonstrating high hormone dependency of these tumors (Fig 5A), although the majority of the tumors in this model exhibit the Luminal B subtype (Fig 3). (Sham control group shows stagnated growth, possibly due to increased apoptosis and senescence that counter moderate rates of proliferation.) However, after a long latency of approximately 6 months, two of the three tumors in ovariectomized rats reappeared, mimicking human ER+/PR+ breast cancers that eventually return after successful treatment and remission for years.

**Figure 5.**
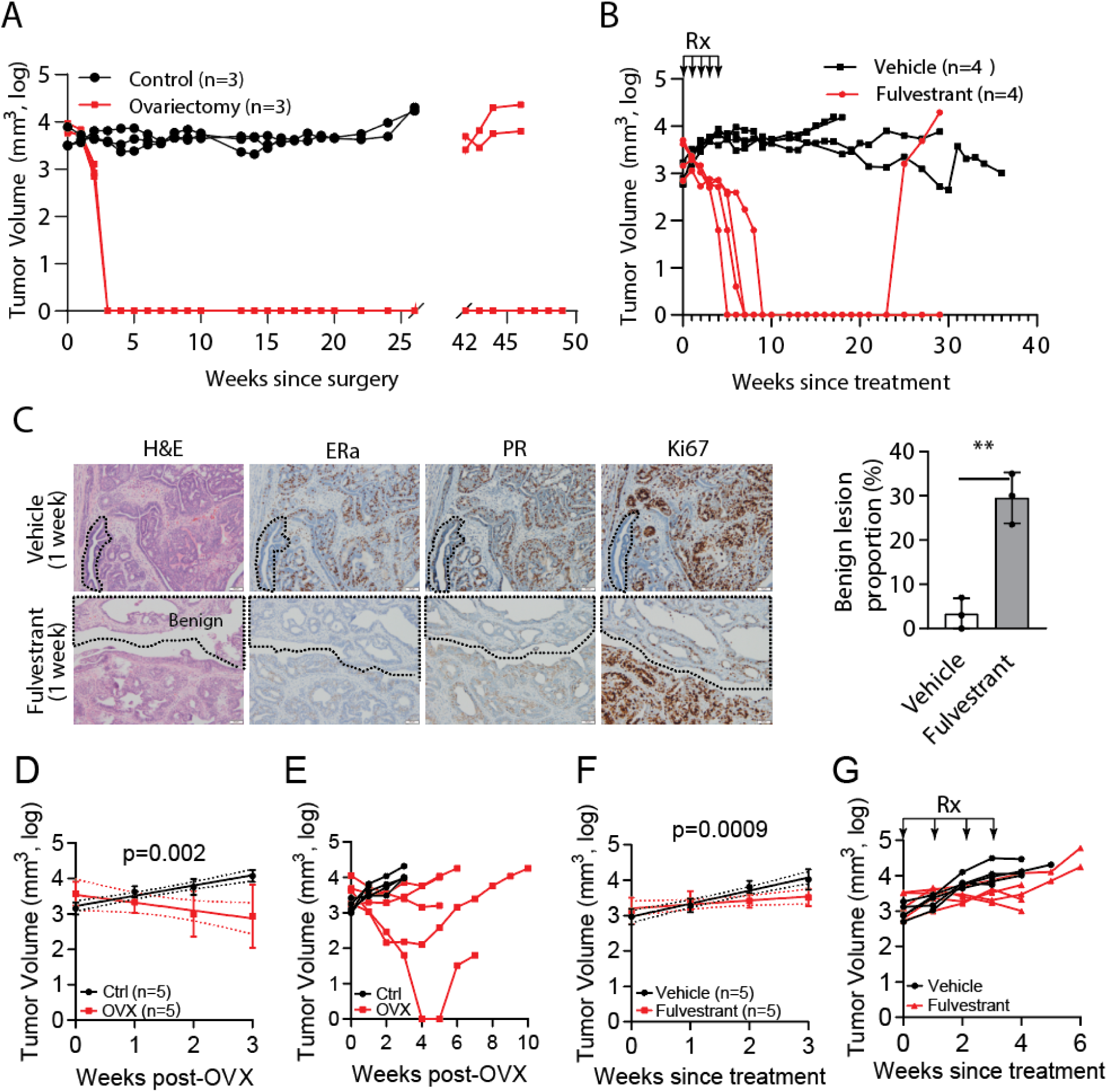
**Tumors induced by *Nf1***^Indel^ **are endocrine-dependent, but those induced by Nf1indel/Tp53indel are endocrine-resistant.** (A and B) Tumor growth curves of rats bearing Nf1 indel mammary tumors treated with ovariectomy (A) or fulvestrant (125mg/kg; 5 weekly s.c. injection). (C) Representative H&E and immunohistochemistry (IHC) staining for ER_α_, PR, and Ki-67 of rat mammary tumors treated with vehicle (top row) or fulvestrant (bottom row) for one week. Dashed lines delineate tumor regions showing therapy-induced regression, characterized by broad, smoothly contoured ducts/lobules, flattened to low-cuboidal epithelium, prominent lumina, minimal nuclear atypia, reduced ER_α_ and PR expression, and low proliferation (Ki-67 staining). (D) Quantification of the proportion of tumor area occupied by differentiated ducts/lobules in vehicle-versus fulvestrant-treated tumors (**P < 0.01**, Student’s t-test). Data shown as mean ± SD, n = 3 per group. (D-G) Tumor growth curves of rats bearing *Nf1*^Indel^ + *Tp53*^Indel^ tumors treated with ovariectomy or fulvestrant (125mg/kg; 4 weekly s.c. injection). Early time points show a modest endocrine response (D and F), but resistance develops rapidly (E and G).

Besides estrogen deprivation, another endocrine therapeutic approach is a class of selective estrogen receptor down-regulators (SERDs), which bind to ER with high affinity and impair ER dimerization and nuclear localization while promoting ubiquitination and degradation. Therefore, we next tested the response of *Nf1* Indel tumors to fulvestrant, a widely used SERD. A 5-week course of fulvestrant treatment caused rapid and full regression of all four tumors (Figure 5B), confirming the tumor dependence on ER, as revealed by ovariectomy. Even without resuming fulvestrant, three of the four rats stayed in remission for the entire observation window of 8 months while the fourth one relapsed at five months following the cessation of treatment. Together, these two endocrine treatment regimens show that *Nf1*^Indel^-induced tumors highly depend on estrogen signaling for continued growth, but in the absence of substantial estrogen signaling such as under estrogen deprivation or fulvestrant treatment, the residual tumors can regain growth, mimicking the clinical behavior of a significant proportion of human ER+/PR+ breast cancer.

Consistent with its high endocrine response, at day 7 of fulvestrant treatment, the tumor mass has morphed into broad, smoothly contoured ducts lined by flattened to low-cuboidal epithelial cells. These areas displayed minimal cytologic atypia, prominent lumina, and increased fibrous stroma, features consistent with therapy-induced regression. Correspondingly, these transfigured regions exhibited near-complete loss of ER_α_ and PR expression and a substantially reduced Ki67 proliferation index (Figure 5C).

Having demonstrated a model that mimics endocrine therapy-sensitive human breast cancer, we next asked whether one of our rat models mimics endocrine therapy-resistant human breast cancer. A *Tp53*^Indel^ mutation in cancer disrupts normal control of cell cycle arrest and cell apoptosis as well as senescence, all of which are critical in tumor growth. A *TP53* mutation in patients has been associated with endocrine resistance in multiple clinical studies ^25,26^. Therefore, we next tested whether mammary tumors induced by *Nf1*^Indel^/*Tp53*^Indel^ may depend less on estrogen signaling. Ovariectomy of the rats bearing *Nf1*^Indel^/*Tp53*^Indel^ tumors caused detectable tumor shrinkage by week 3 (p<0.01), but the response was very modest—the tumor mass in four of the five rats never fully regressed (Fig 5D). Importantly, by week 4 to 5, all tumors resumed growth (Fig 5E). In accordance with this finding, fulvestrant treatment of rats bearing these tumors caused a brief regression (p<0.05 at week 3 compared to the control cohort) (Fig 5F) but failed to produce a sustained and significant benefit compared to the vehicle-treated control rats (Fig 5E). Together these two endocrine treatment experiments indicate that these *Nf1*/*Tp53*-double-mutated tumors only partly depend on estrogen signaling and gain resistance quickly, closely mimicking the ER+/PR+/Ki67^high^ human breast cancers that are resistant to endocrine therapy.

### Transcriptome changes in rat models in response to endocrine treatment reveal critical transcriptome changes in human ER+ tumors treated with endocrine therapies

Having tested their endocrine response, we next investigated their transcriptome changes and their relevance of estrogen and endocrine therapy-induced transcriptome changes in human breast cancer. A total of 486 genes were differentially expressed in either model following fulvestrant treatment (7 days for the *Nf1*^Indel^ model and 5 weeks for the *Nf1*^Indel^/*Tp53*^Indel^ model; Fig 6A). Many of these genes overlap with the ER-responsive genes previously reported in cultured human ER+ breast cancer cells ^27^. Examples include PGR^28^, AREG ^29^, SGK3 ^30,31^, KLF10/TIEG1 ^32^, STC2 ^33^, TSKU ^34^, DHFR ^35^, PFKFB3 ^36^, PHB^37^, AGPAT5 ^38^, TIPARP ^39^, WNT5A ^40^, and CXCL3 ^41^. Notably, some of the expression pattens are shared across the two models while others are model-specific.

**Figure 6.**
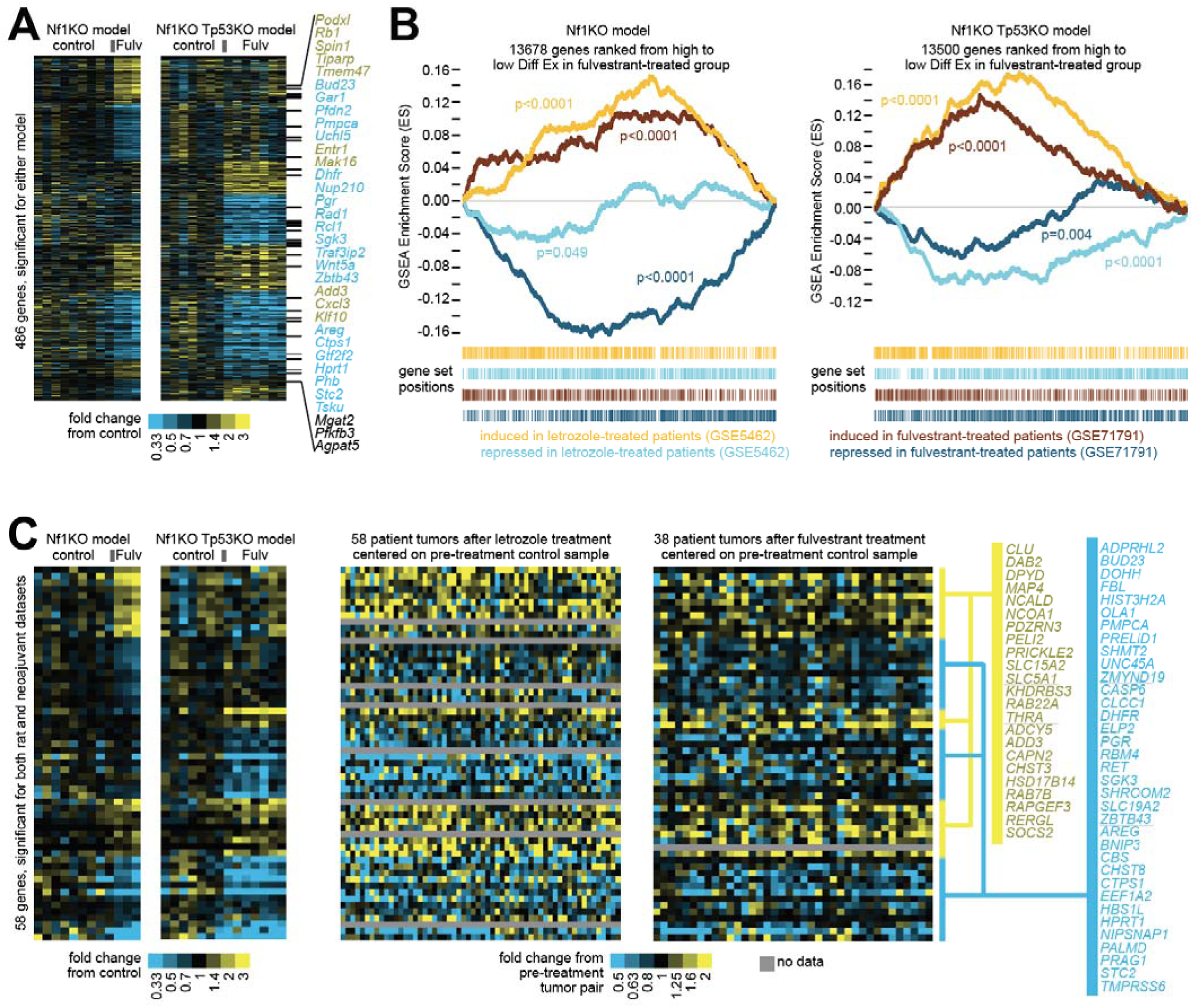
Genes differentially expressed with fulvestrant treatment in rat models have orthologs similarly affected in human breast tumors treated with endocrine therapy. (A) For Nf1KO and Nf1KO/Tp53KO rat models, genes differentially expressed with fulvestrant treatment in either model (p<0.01, t-test). Genes are sorted according to their differential patterns being specific to one model or common to both models. Genes listed to the side are those for which human orthologs were induced by estrogen in vitro [PMID: 16606439]. (B) For each rat model, genes profiled were ranked by high to low differential expression (Diff Ex) in fulvestrant-treated versus control groups. GSEA plots evaluated enrichment with the fulvestrant-treated tumors for gene orthologs differentially expressed in human tumors from patients treated with endocrine therapy in the neoadjuvant setting. For the two human datasets considered (GSE71791, fulvestrant; GSE5462, letrozole), tumors from patients both pre- and post-treatment were profiled for gene expression, and the top differential genes (either higher or lo post-treatment, using p<0.01 by paired t-test) were examined in the context of the rat gene orthologs ranked according to the rat tumor data. Genes induced by neoadjuvant treatment in the human setting are significantly enriched in fulvestrant-treated rat tumors (indicated by positive GSEA plot curves); genes repressed in the human setting are anti-enriched in fulvestrant-treated rat tumors (negative GSEA curves). P-values by GSEA-based permutation testing. (C) From the genes in part a, the top subset that were also differentially expressed in the breast cancer patient neoadjuvant setting for either human dataset (p<0.01, paired t-test). Gene ordering is the same across heat map panels.

We next asked whether these transcriptome changes recapitulate those observed in patients following endocrine therapy. Using gene set enrichment analysis (GSEA), we compared the transcriptome changes in rats versus patients. As shown in Fig 6B, ranking all profiled human gene orthologs from the *Nf1*^Indel^ model (from high to low differential expression and excluding those with zero values) revealed significant enrichment of genes induced in human breast cancer after neoadjuvant treatment with either letrozole (GSE5462) ^42^ or fulvestrant (GSE71791) ^43^ and significant depletion of the corresponding repressed genes. A similar pattern was observed when comparing the *Nf1*^Indel^/*Tp53*^Indel^ model data to the patient data. Of note, while our two rat models are distinct in endocrine response, patients in these two trials were accrued without prior knowledge of endocrine sensitivity, and thus each dataset included both responsive and resistant tumors.

We further visualized the similarity of endocrine responses between our two rat models and patients using heatmaps. A set of 58 genes were differentially altered in one or both rat models and also altered in either human dataset (Fig 6C). Again, many of these genes are known to be regulated by estrogen, including AREG ^29^, SGK3 ^30,31^, STC2 ^33^, DHFR ^35^, as also shown in Fig 6A. Importantly, both over-represented and depleted gene clusters in the two rat models were also readily observed in the two huma datasets. Collectively, these comparisons demonstrate that the transcriptome changes in our two rat models show strong concordance with those in patient tumors.

### Somatic genome editing-induced mammary models in rats show the microenvironment diversity of ER+/PR+ human breast cancer

Tumor microenvironment, especially the immune microenvironment, is critical in shaping tumor formation and progression as well as therapeutic response/resistance. However, due to the scarcity of mouse models of ER+ breast cancer, this disease is most commonly studied in immunodeficient hosts. As a result, the contributions of tumor microenvironment, especially the immune microenvironment, could not be properly investigated. Having developed a panel of ER+/PR models, we decided to use scRNA sequencing to profile their microenvironment. UMAP analysis across all rat models revealed a broad variety of non-epithelial cell types including major immune cell populations, fibroblasts, pericytes and endothelial cells (Fig 7A). Focusing on immune cells, we observed marked heterogeneity in both adaptive immune cells and myeloid cells across different models. As shown in Fig 7B & C, adaptive immune cells were enriched in the tumors initiated by *Pik3ca^H1047R^*alone or *Tp53*^Indel^ alone but are depleted in *Nf1^indel^* tumors and in models driven by more than one genetic lesion. Expansion of myeloid cells was associated with decrease of adaptive immune cells across the models. Interestingly, we observed a dichotomy between enrichment of macrophages and plasmacytoid dendritic cells in *Nf1^Indel^ and Nf1^Indel^/Tp53^Indel^*tumors and enrichment of neutrophils in *Tp53^Indel^/Pik3ca^H1047R^* and *Nf1^Indel^/Tp53^Indel^/Pik3ca^H1047R^* tumors. This macrophage-neutrophil dichotomy parallels our previous findings in murine TNBC transplantable models and human patients ^44^. Furthermore, the enrichment of neutrophils in *Tp53^Indel^/Pik3ca^H1047R^* and *Nf1^Indel^/Tp53^Indel^/Pik3ca^H1047R^* tumors may reflect enhanced mTOR signaling (“group 4” in Fig 3), consistent with the role of mTOR in promoting systemic neutrophil accumulation and tumor infiltration ^45^. Together, these data demonstrate that rat models provide a powerful toolkit to investigate the heterogeneous tumor microenvironment of ER+ tumors.

**Figure 7.**
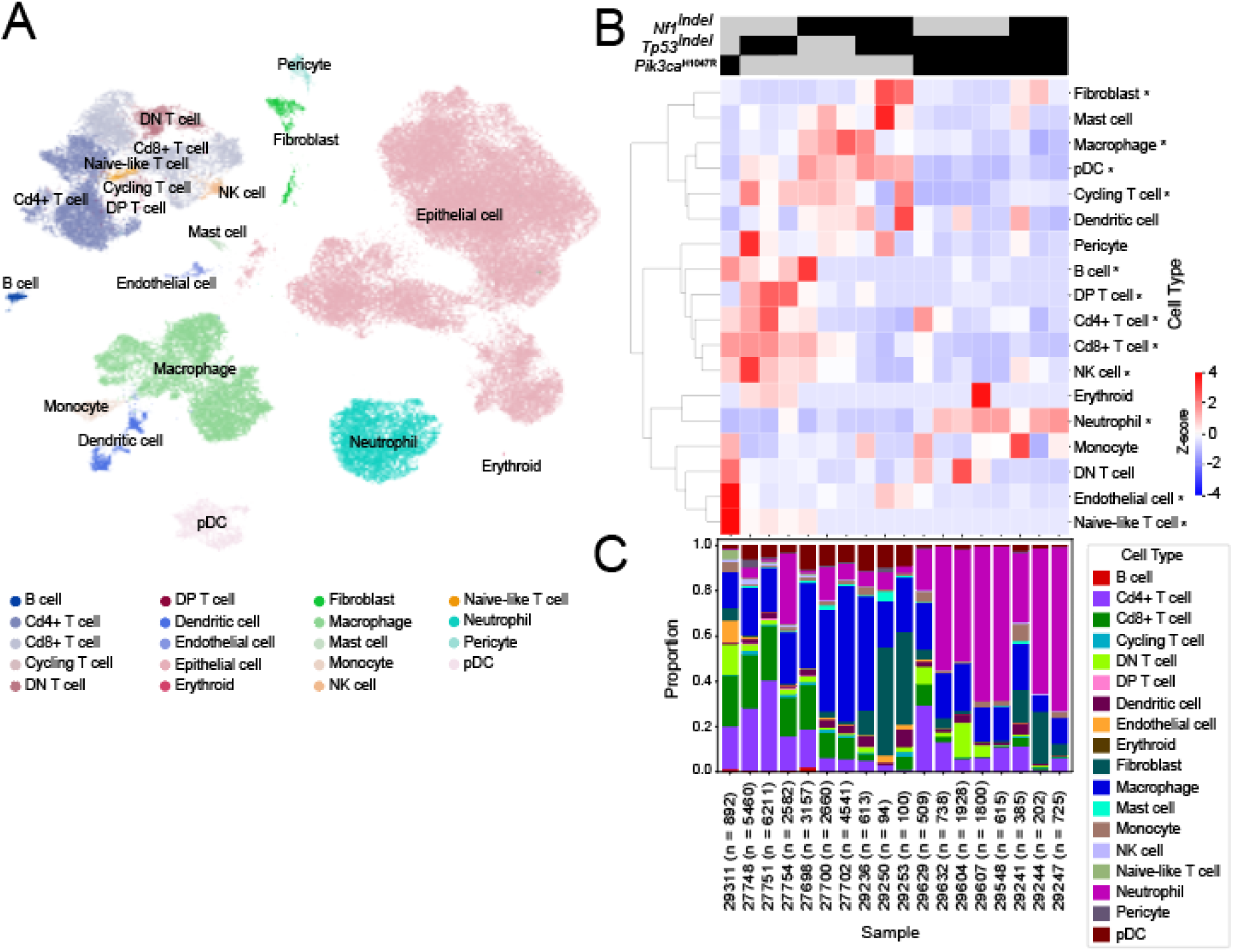
Somatic genome-editing-induced mammary tumors in rats exhibit distinct tumor microenvironments depending on oncogenic drivers. (A) UMAP visualization of 78,221 cells analyzed by scRNA-seq and integrated across 18 rat primary breast tumors. Cells were integrated using scVI/scANVI and embedded in two dimensions using UMAP. Clusters represent transcriptionally distinct cell populations identified via Leiden clustering. Cell types were annotated using a consensus approach that combined reference-based methods (CellTypist and scANVI trained on the Tabula Muris Senis dataset) with marker-based annotation using ScType, leveraging curated gene sets from CellMarker2, PanglaoDB, and the ScType database. (B) Heatmap of z-score–normalized proportions of non-epithelial cell types across rat model scRNA-seq samples. Rows represent annotated cell types, and columns correspond to individual samples. Proportions were calculated as the fraction of total non-epithelial cells per sample assigned to each cell type, followed by row-wise z-score normalization to highlight relative abundance across samples. The blue–white–red (BWR) color scale indicates below-average (blue), average (white), and above-average (red) relative abundance of each cell type. (C) Stacked bar plot of non-epithelial cell type composition across individual rat model scRNA-seq samples. Each bar represents a sample, with colors corresponding to distinct immune and stromal cell types identified by single-cell RNA sequencing. The height of each segment reflects the proportion of that cell type among all non-epithelial cells within the sample.

## Discussion

We established a somatic CRISPR genome-editing platform in rats that enables rapid, tissue-specific introduction of oncogenic alterations without germline manipulation. We applied this approach for multiplex introduction of patient-relevant alterations into adult mammary epithelium and generated multiple models of sporadic breast tumorigenesis in an immunocompetent setting. Beyond breast cancer, the same framework is extensible to other genes and tissues, offering a scalable strategy for building rat models of human cancers that remain underrepresented in mice.

A central finding is the species-specific divergence in mammary tumor biology under identical genetic drivers. Whereas common mouse approaches frequently yield ER-negative, metaplastic tumors with prominent squamous features, a rare sub-type in breast cancer patients comprising less than 0.3% of breast cancer cases ^46^, our rat models consistently generate ER-positive ductal lesions with little to no metaplasia. This pattern holds across multiple genetic contexts—including *Kras*^G12D^, *Pik3ca*^H1047R^, and combined *Pik3ca^H^*^1047R^/*Tp53*^Indel^—and underscores intrinsic differences in lineage plasticity and/or tissue context that favor ER-positive tumorigenesis in rats. These data indicate that rats uniquely recapitulate the predominant human phenotype—NOS ductal carcinoma that is ER+/PR+—and thus provide an urgently needed complement to mouse models for mechanistic studies of hormone-driven disease and for testing new therapeutics in prevention and treatment.

Importantly, the platform yields models that bracket clinically relevant behavior to endocrine treatment. Tumors initiated by *Nf1^Indel^* display robust estrogen dependence and durable regressions after ovariectomy or fulvestrant, with occasional late relapse—mirroring endocrine-sensitive human disease that can recur after remission. In contrast, *Nf1*^Indel^/*Tp53*^Indel^ tumors show only partial, transient regressions and early regrowth, modeling endocrine-refractory disease frequently seen in patient breast tumors with TP53 alterations. These complementary phenotypes provide tractable systems to dissect how NF1 loss confers endocrine dependence, how TP53 disruption erodes it, and how residual disease adapts under estrogen pathway pressure. The transcriptomic concordance we observe between rat tumors under endocrine therapy and neoadjuvant patient datasets further supports their translational fidelity and suggests immediate value for testing SERDs, CDK4/6 inhibitors, and rational combinations guided by pathway signatures. These models may also prove useful for studies of dormancy that is common in ER+ breast cancer and accounts for late recurrence in some patients.

At the tumor-microenvironment level, single-cell profiling reveals heterogeneous immune ecosystems across genotypes, providing a rich resource for exploring tumor-immune microenvironment crosstalk. ER+ breast cancers are generally thought to be immunologically cold and not responsive to immunotherapies due to a lack of high mutational burden and neoantigens ^47^. As a result, the roles of immune cells in ER+ tumor progression have not been extensively studied. Consistent with the general belief, adaptive immune cells are low in most of our models. However, our data revealed an unexpected enrichment of heterogeneous myeloid cells including neutrophils and macrophages. Both neutrophils and macrophages can blunt anti-tumor immunity ^48,49^. Their accumulation raises the possibility that the cold microenvironment may not be due to lack of mutations or neoantigens but the active recruitment of suppressive myeloid cells. Interestingly, there appears to be a dichotomy between neutrophil and macrophage enrichment across different models, suggesting mutually exclusive, parallel pathways toward a cold tumor. Identification of the underlying mechanisms may lead to novel therapeutic strategies to reprogram the tumor microenvironment and unleash anti-tumor immunity.

These rat models also expand our toolkits to study different stages of breast cancer. The *Nf1*^Indel^ tumors with characteristics of DCIS as well as the presence of the DCIS stage in other rat models in this study provide an opportunity to study DCIS progression to invasive cancer. DCIS is detected in approximately 50,000 cases a year in the U.S and is the non-obligatory immediate precursor to invasive cancer, but studying this progression has been hampered in mouse models since DCIS is rarely reported in GEM models ^1^. On the other hand, the *Pik3ca*^H1047R^ model presents a new opportunity to investigate fibroadenoma genesis and progression from this benign lesion type to cancer. In addition, as we generate more tumors using this platform and characterize their distant metastases, we may be able to offer new models for studying ER+ breast cancer metastases, an area that is poorly modeled by mice ^1^.

Limitations include the current focus on luminal tumors driven by select alterations; while representative of the majority of human disease, broader coverage (e.g., ESR1 ligand-binding-domain mutations, MAP3K1, GATA3, and chromatin regulators) will refine genotype–phenotype maps. Although we minimize confounders by editing defined alleles, long latencies in some settings imply accrual of secondary events; future barcoding/lineage tracing and whole-genome interrogation can parse necessary cancer-permissive changes. Finally, head-to-head immune and stromal benchmarking across rat, mouse, and human tissues will help pinpoint the non-cell-autonomous determinants that bias lineage outcomes across species.

In summary, rat somatic genome editing closes a long-standing species gap for ER-positive breast cancer, aligning histology, endocrine dependence, and treatment-induced transcriptome remodeling with clinical observations. These models should accelerate mechanism-based prevention studies, illuminate resistance trajectories under modern endocrine regimens, and enable rational design of combination strategies that jointly target tumor-intrinsic pathways and the immune–stromal milieu.

## Materials and Methods Animals

LE-ROSA26^tm1(LSL-Cas9)Ottc^ and Wistar-Tg(CAG-Ncre)81Jmsk rats were purchased from Rat Resource & Research Center (RRRC#: 00833 and 301, respectively). Rats constitutively expressing Cas9 were generated by breeding LE-ROSA26^tm1(LSL-Cas9)Ottc^ and Wistar-Tg(CAG-Ncre)81Jmsk rats and genotyped according to the protocol provided by RRRC. All rats were kept on Standard Rodent Diet (5V5R) (Lab Diet Picolab Select Rodent 50 IF/9F(extruded)). *Gt(ROSA)26Sortm1.1(CAG-cas9*,-EGFP)Fezh*] mice were purchased from The Jackson Laboratory (Bar Harbor, ME). All mice were kept on 2920X Teklad Global Extruded Rodent Diet (Soy Protein-Free; Harlan Laboratories, Indianapolis, IN). Only female rats and mice were used in this study. All procedures using rat were performed in compliance with an Institutional Animal Care and Use Committee–approved animal protocol.

### Plasmids

Packaging plasmids pAAV2/9n and pAdDeltaF6 were obtained from Addgene (plasmids #112865 and #112867). AAV-GFP was purchased from Addgene (Plasmid #49055). AAV-P and AAV-K for editing mouse *Pik3ca*^H1047R^ and KrasG12D have been reported before ^6^. An intermediate AAV-rNf1-L construct was constructed by removing HDR donor region between HindIII and SacI of AAV-KPL (Addgene #60224) and replaced the NdeI and KpnI region with a synthesized fragment (5’CATATGCTTACCGTAACTTGAAAGTATTTCGATTTCTTGGCTTTATATATCTTGTGGAAAGGACGAAACACCACTAATCCTTAACTACCCAAGTTTTAGAGCTAGAAATAGCAAGTTAAAATAAGGCTAGTCCGTTATCAACTTGAAAAAGTGGCACCGAGTCGGTGCTTTTTTGGTACC) containing rat sgRNA targeting rat *Nf1* gene. Another intermediate construct Lenti-CRISPR v2-*Tp53*gRNA was constructed by inserting a fragment generated from two annealed oligos (5’ CACCGcatcgagctccctctgagtc and 5’ AAACgactcagagggagctcgatgC) containing a gRNA targeting rat *Tp53* gene into the BsmBI site of Lenti-CRISPR v2. The region between KpnI and NheI of AAV-rNf1-L was replaced by the fragment between KpnI and NheI from Lenti-CRISPR V2-rP53gRNA resulting AAV-NT. The AAV-T was constructed by removing the fragment between MluI and KpnI of AAV-NT.

The AAV-N was constructed by removing the region between KpnI and NheI of AAV-NT. To make the rat *Pik3ca*^H1047R^ editing, the AAV-rP was constructed by multiple steps. First, the gRNA targeting mouse *Pik3ca* gene in the AAV-P ^6^ was converted into rat gNRA targeting rat *Pik3ca* gene through site-directed mutagenesis using primers: 5’ ATGAATGAcGCACATCATGGGTTTTAGAGC and 5’ CGGTGTTTCGTCCTTTCCAC. The region between NdeI and NheI from the resulting vector was used to replace the region between NdeI and NheI of AAV-NT resulting an intermediate vector AAV-rPgRNA. To create the HDR donor sequence for rat *Pik3ca*^H1047R^ editing, the rat genomic DNA from a Cas9/Cre double transgenic rat was used as template for PCR-cloning an 813bp fragment spanning the editing region into the PCR2.1 vector. The primers used are 5’ TACTGAACTCCAATGTTCC and 5’ CCATCATTTGTATGTATTTTG. The wild type HDR donor region in this vector was converted into H1047R editing version by site-directed mutagenesis using the primers: 5’ ATGAACGATGCCCGCCACGGTGGCTGGACAACAAAAATGGACTGGATC and TTGCTTTGTGAAATACTCCAG. The mutated HDR donor was PCR amplified (primers: 5’-tccacttctcaggtaaccactactgaactccaatgttcc and 5’-tgcggccgctcggtccgcacccatcatttgtatgtattttg) and In-Fusion-inserted into the PmlI site of AAV-rPgRNA intermediate vector to generate the final AAV-rP vector. The AAV-rPNT vector was constructed by In-Fusion-inserting the sgRNA region from AAV-NT (PCR using primers: 5’ GTGCTTTTTTGCTAGCGGCCGCACGCGTGAG and 5’ CTCCGCTTCCGCTAGC) into AAV-rP vector at the NheI site. The AAV-rPT was generated by replacing the NheI region of AAV-rPNT with a PCR fragment using AAV-rT as template and primers 5’ GTGCTTTTTTGCTAGCGAGGGCCTATTTCCCATG AND 5’ CTCCGCTTCCGCTAGC through In-Fusion cloning. To generate AAV-rK, the region between HindIII and and PmlI of mouse *Kras*^G12D^ editing vector AAV-K ^6^ was first replaced by an 825bp PCR fragment (rat genomic DNA as template and primers: 5’ TTTTTTGTGTAAGCTTAGCTAGCTGTCAACAAGCTC AND 5’ CCGCTCGGTCCGCACGTGCAACAACACAATAACAAACAAACCC) through In-fusion cloning resulting an intermediate vector #1. A site-directed mutagenesis was subsequently conducted to replace the mouse gRNA with the rat gRNA in the intermediate vector #1 resulting an intermediate vector #2 using primers: 5’ TGTATCGTCAGTTTTAGAGCTAGAAATAGCAAG and 5’ GCTAATTCAGGGTGTTTCGTCCTTTC. To generate a rat *Kras*^G12D^ HDR donor sequence, fusion PCR was carried out using the 825bp of wild type HDR donor sequence as template and primers 5’ TTTTTTGTGTAAGCTTAGCTAGCTGTCAACAAGCTC and 5’ GCTGGATTGTTAATGCACTCTTGCCTACGCCATCAGCTCCAACTACC for one fragment, and 5’ AGTGCATTAACAATCCAGCTAATTCAGAATCACTTTG and CCGCTCGGTCCGCACGTGCAACAACACAATAACAAACAAACCC for the other overlapping fragment; and primers 5’ TTTTTTGTGTAAGCTTAGCTAGCTGTCAACAAGCTC and 5’ CCGCTCGGTCCGCACGTGCAACAACACAATAACAAACAAACCC for the full-length HDR donor fragment. The resulting full-length rat *Kras*^G12D^ HDR donor was then used to replace the wild type HDR donor in the intermediate vector #2 through In-fusion cloning resulting the final construct AAV-rK. To generate mouse *Pik3ca*^H1047R^ and *Tp53*^Indel^ editing vector AAV-mPT, a synthesized DNA fragment containing gRNA targeting mouse *Pik3ca* (catatgcttaccgtaacttgaaagtatttcgatttcttggctttatatatcttgtggaaaggacgaaacaccgATGAATGATGCACATCATGGgttttagagctagaaatagcaagttaaaataaggctagtccgttatcaacttgaaaaagtggcaccgagtcggtgcttttttgGATCc) was used to replace the region between NdeI and BamH1 of AAV-K ^6^ resulting intermediate vector #3. The fragment between MluI and BamHI in intermediate vector #3 was used to replace the region between MluI and BamHI of AAV-KPL^50^ after its XbaI and AgeI region was removed resulting an intermediate vector #4. The region between ApaI and SacI of intermediate vector #4 was then replaced by the region between ApaI and SacI of AAV-P ^6^ resulting the final construct AAV-mPT.

### Virus production

Virus production has been previously described ^6^. In brief, the jetPRIME transfection reagent (Polyplus transfection, 114-07) was used for plasmid transfection, the AAVpro Purification kit (Takara #6675) was used for virus extraction/purification/concentration, and the AAVpro Titration kit (Takara #6233) was used for titer determination following the manufacturer_’_s instructions.

### Intraductal injection

Intraductal injection of virus has been previously described ^24,51^. One mammary gland (usually the right #4 gland) of each rat was injected.

### Mammary gland fluorescent signal imaging

Mammary glands were collected from euthanized rats and imaged immediately under a fluorescent stereomicroscope (Leica MZ 16 F) through its matched software (LAS 3.8).

### Preparation of single-cell suspensions from mammary glands and flow cytometry analysis

Preparation of single-cell suspensions from mammary glands and flow cytometry analysis of GFP+ cells have been described previously ^52^. For tumor immune cell profiling, rat tumor samples were minced into small pieces and digested with 1.5 mM collagenase I (17018-029, Gibco) and 1_μ_g/ml DNase I (DN25, Sigma-Aldrich) at 37°C for 1 hour with continuous rotation. The digested samples were passed through a 70μm cell strainer and then washed with PBS containing 2% fetal bovine serum (FBS) before proceeding to staining. The following anti-rat antibodies were used for rat cell staining: CD45-PB (202226, BioLegend), CD11b-Percp/Cy5.5 (201820, BioLegend), His48-FITC (11-0570-82, Invitrogen), CD43-PE-Cy7 (202816, BioLegend), CD3-PE (201412, BioLegend), CD4-APC/Cy7 (201518, BioLegend), CD8-AF647 (201710, BioLegend). CD45R-BV711 (743594, BD Biosciences). CD161-PE (205604, BioLegend), CD3-FITC (11-0030-85, Invitrogen), CD103-AF647 (205509, Biolegend). The gating strategies for rat cells are: Neutrophil: CD45^+^CD11b^+^His48^+^CD43^+^; Monocyte: CD45^+^CD11b^+^His^-^CD43^+^; Macrophage: CD45^+^CD11b^+^His48^med^CD43^med^; B cell: CD45^+^CD11b^-^CD3^-^CD45R^+^; Total T cell: CD45^+^CD11b^+^CD45R^-^CD3^+^; CD4 T cell: CD45^+^CD11b^+^ CD45R^-^CD3^+^CD4^+^; CD8 T cell: CD45^+^CD11b^+^CD45R^-^CD3^+^CD8^+^. NK cell: CD45^+^CD161^+^CD3^-^; NKT cell: CD45^+^CD161^+^CD3^+^; DC: CD45^+^CD103^+^.

### Amplicon sequencing

Genomic DNA extraction has been previously reported ^6^. Purified tumor genomic DNA was used as template of PCR for amplifying the edited region using following primers: 5’ CTAAAGTTGCTCCCTAGTG and 5’ CAACAGATCTATTAATACAC for *Nf1,* editing, 5’ GACAAACTATGCATCCATAC and 5’ GGGATAGCACCTCAGGGGC for *Tp53* editing, and 5’ AGCCTTAGACAAAACTGAGC and 5’ CCCAGCTGCCATCTCAGTTC for *Pik3ca* editing. The PCR product was purified using QIAquick® PCR Gel Cleanup kit (QUIAGEN Cat. No. 28506) and sequenced by Amplicon Sequencing (Azenta Life Science).

### Drug treatment

Fulvestrant (Selleckchem #S1191) was prepared as follows: 80mg fulvestrant was dissolved in 80_μ_l DMSO followed by mixing with 1520_μ_l corn oil resulting in a final concentration of 50mg/ml. A dose of 2.5_μ_l/g body weight (125mg/kg body weight) was administered through subcutaneous injection weekly.

The same mixture of DMSO and corn oil was used as a vehicle and administered with 2.5_μ_l/g body weight weekly.

### Ovariectomy

Under isoflurane anesthesia, artificial tears was placed in each eye. The surgical site was clipped and then any additional hair will be removed with depilatory cream (such as Nair®). The surgical site was then prepared aseptically with three alternating preps of betadine of chlorhexidine scrub and alcohol. Sterile technique was used, by using autoclaved instruments, sterile surgical gloves, sterile drapes and aseptic technique. A local block of lidocaine bupivacaine was applied prior to all incisions. The reproductive tract is exposed through a longitudinal incision. The ovaries are isolated, ligated with sterile vicryl absorbable suture material, and excised. The reproductive tract is returned to the peritoneum and the wound-closed with sterile vicryl absorbable suture for inner layer and wound clips for skin. Buprenorphine (0.05 mg/kg) and meloxicam (2 mg/kg) were administered subcutaneously 1 hour before surgery. Buprenorphine was repeated 12 hours after surgery, and meloxicam was administered once daily for 3 days postoperatively.

### Tissue processing and H&E staining

Tissue processing and hematoxylin and eosin (H&E) staining have been previously described ^52^. Antibodies used are: Estrogen Receptor alpha Monoclonal Antibody (6F11) (Invitrogen, MA1-80216, 1:200), Progesterone Receptor Monoclonal Antibody (hPRa 7) (Invitrogen, MA5-12658, 1:200) or Progesterone Receptor pAb (Abclonal, A0321, 1:800), BD Pharmingen™ Purified Anti-Ki-67 (Clone B56 (RUO)) (BD Bioscences, 5506609, 1:200), p63 Antibody (4A4) (SCBT, sc-8431, 1:200). The proportion of ER, PR and Ki67 were semi-estimated. The staining intensity of ER and PR were scored as 1, 2, and 3 for weak, medium, and strong staining.

### RNA-Seq alignment and quantification

Tumors used for RNA-Seq were snap-frozen in liquid nitrogen and subsequently stored at −80 °C before RNA extraction. Total RNA was extracted using the Direct-zol RNA Miniprep Plus Kit (Zymo Research, R2072) according to the manufacturer’s instructions. The purified RNA samples were then submitted to Novogene Corporation Inc. (Sacramento, CA, USA) for RNA-Seq. Quality control was performed using the Agilent 5400 Fragment Analyzer System to evaluate RNA quantity, purity, and integrity. RNA libraries were prepared using poly(A) enrichment, and sequencing was carried out on the Illumina NovaSeq X Plus platform with a 150 bp paired-end strategy.

RNA-seq reads were aligned to the rat reference genome (Rnor_6.0, NCBI, along with refGene.gtf file for gene annotations) or to the mouse reference genome (mm10, ENnemble) using Bowtie2, and gene-level expression quantification was performed using RSEM ^53^. RSEM estimated transcript abundance and generated gene-level expected counts and RPM values. The resulting expression data were log-transformed and quantile normalized ^54^ to correct for differences in sequencing depth and RNA composition across samples by organism type. All tools were run with default parameters unless otherwise specified.

Differentially expressed genes across rat model categories were identified using one-way analysis of variance (ANOVA) on log2-transformed values. Genes with an adjusted False Discovery Rate (FDR) < 1% (Storey-Tibshirani correction for multiple testing ^55^) were used for unsupervised clustering using Cluster 3.0 ^56^ and JavaTreeView ^57^. Rat tumor expression profiles were each assigned an expression-based intrinsic based on the Hoadley et al. human expression dataset ^58^, using a previously described analytical approach ^59^. Briefly, for each gene common to our rat tumor profiling dataset 9 using human orthologs) and the Hoadley dataset, we computed the mean centroid of the four major Hoadley tumor subtypes (Luminal A, Luminal B, Basal-like, HER2+) and centered each group average on the centroid. With our rat tumor dataset, we centered the log2 expression values to standard deviations from the median across tumor profiles. We then took the Pearson’s correlation (using all genes common to both data sets) between the Hoadley centered averages and the differential expression values of each rat tumor profile, and the subtype with centroid having the highest correlation was assigned to the sample profile. If no intrinsic subtype has a Pearson’s correlation with significance of p<0.05, the subtype was considered as indeterminant. Pathway enrichment for selected gene clusters was carried out using SigTerms ^60^. For TCGA breast cancer datasets, mutation status was previously compiled ^61^

Significance of gene set enrichment involving gene expression profiles from the fulvestrant-treated rat tumors was determined using the Gene Set Enrichment Analysis (GSEA) software ^62^, with “classic” Enrichment Score metric and 10000 randomly generated gene sets. Group comparisons were based on t-tests on log2-transformed data. Genes in the given rat tumor dataset that both had non-zero values across all tumors and a human ortholog were used to rank the genes from greatest induction by fulvestrant (by t-statistic) to greatest repression.

### scRNA-Seq

Tumor tissue processing methods were described in the flow cytometrypart. Sorted live cells were used to construct single cell gene expression library according to the Chromium Single Cell Gene Expression 3v3.1 kit (10x Genomics). The scRNASeq was conducted by Illumina X platform at BCM RNA Profiling Core.

Raw FASTQ files were preprocessed using 10x Genomics Cell Ranger v8.0.0 [28091601] to generate gene-barcode count matrices aligned to the appropriate reference genome: mRatBN7.2 (Ensembl release 112) for rat samples and GRCm39 (Ensembl release 112) for mouse samples. Downstream analysis was performed using Seurat (v5.1.0) [37231261] and Scanpy (v1.10.3) [29409532] in R (4.3.3) and Python (3.11.9), where samples were first individually processed then integrated.

Initial quality control and filtering were conducted using Seurat. Cells were retained if they expressed at least 200 genes and had less than 10% mitochondrial RNA content. Genes were retained if detected in at least three cells. To exclude outliers, cells with feature counts below the 0.5th percentile or above the 98th percentile within each sample were removed. Putative doublets were identified and excluded from analysis using DoubletFinder [30954475].

Normalization was performed using SCTransform [31870423]. Dimensionality reduction was carried out using principal component analysis (PCA), retaining the top 30 components. A shared nearest neighbor (SNN) graph was constructed using Seurat’s *FindNeighbors* on the PCA-reduced space, and cells were embedded in two dimensions using Uniform Manifold Approximation and Projection (UMAP) via *RunUMAP*. Clustering was performed using Seurat’s *FindClusters* function across a range of resolutions (0.4–2.0), with the optimal resolution selected based on visual inspection and Clustree visualization [30010766]. A consensus of multiple approaches was used for cell type annotation: Reference-based annotation was performed using CellTypist [35549406] and scANVI [33491336], both trained on the Tabula Muris Senis (droplet) mouse reference dataset [32669714].

Marker-based annotation was carried out using ScType [35273156], incorporating curated markers from the CellMarker2, PanglaoDB, and ScType databases [35273156, 36300619, 30951143]. After individual sample annotation, datasets were integrated using scVI [30504886] and scANVI [<u>33491336</u>] with the top 6,000 highly variable genes to correct for batch effects and improve cross-sample label consistency. Leiden clustering was performed on the integrated latent space using Scanpy, and cell type labels were refined based on these clusters through manual inspection of differentially expressed marker genes. Integrated data were used for UMAP visualization.

Cell type proportions across experimental conditions were visualized using a z-scored seaborn clustermap, Samples were ordered based on predefined experimental conditions, and hierarchical clustering (Ward’s method) was applied to the cell types (rows). Epithelial cells were excluded, including the ambiguous cell clusters which had mix of epithelial and other cell type markers (stromal or myeloid).

Differential cell type proportion analysis was conducted as follows. Cell type counts were aggregated by sample to construct a pseudobulk matrix. Differences in cell type proportions across groups (models) were assessed using the *propeller* method [36005887] (*speckle* v1.4.0) in R. Proportions were variance-stabilized using the arcsine transformation (transform = "asin"), followed by moderated t-tests with Benjamini-Hochberg false discovery rate (FDR) correction. Cell types with FDR < 0.05 were considered statistically significant.

## Statistics

Each value reported represents the means ± SDs of at least three biological replicates. Student_’_s *t* test (if normally distributed) was used to test the significance of difference between two means. Kaplan-Meier plots were generated by GraphPad software, which uses the log-rank (Mantel-Cox) test. *P* values were two-sided unless otherwise specified. Tumor growth linear regression and slope comparison was performed by GraphPad software, which uses simple linear regression and ANCOVA (Analysis of Covariance) slope comparison test.

## Acknowledgements

This work was financially supported by NIH/NCI (R01 CA271498 to Y. L. and X. Z. and CA271588 to Y.L.), DOD-CDMRP (BC231186 to Y. L. and BC230628 to E. C.). This project was also supported by the Breast Center Pathology Core as part of the SPORE program (P50 CA186784), Cytometry and Cell Sorting Core with funding from CPRIT (RP180672) and NIH (S10 RR024574), Antibody-based Proteomics Core with funding from CPRIT (RP210227) and NIH (S10 OD028648 to S.H.), DDC GEMS core with funding from NIH/NIDDK (DK56338), and resources from the Dan L. Duncan Cancer Center with funding from NIH/NCI (P30 CA125123).

## Author Contributions

**Conceptualization / study design:** W.B. X.Z., Y.L. **Methodology / experiments:** W.B., R.X., L.H., A.Z.B., Y.D., S.H., B.Z. **Data collection:** W.B., R.X. **Data analysis:** W.B., T.L., C.J.C., M.A., C.N.,

C.G., C.K.O., **Writing – original draft:** W.B., Y.L. C.J.C., T.L., M.A., Y.D. **Writing – review & editing:** W.B., X.Z., Y.L. **Supervision:** W.B., X.Z., Y.L. **Funding acquisition:** C.C., J.X., A.S, E.C., X.Z., and Y.L.

## Competing interests

The authors declare that they have no competing interests.

## Data and materials availability

The RNA-seq and single-cell RNA-seq raw sequencing data generated in this study will be deposited in the NCBI Sequence Read Archive (SRA). Processed data, including gene-level counts and cell-level expression matrices with associated metadata, will be available through the NCBI Gene Expression Omnibus (GEO). All other data supporting the findings of this study are available within the article and its Supplementary Information files.

## Additional information

There are 6 supplemental (extended) figures.

## Notes

### Competing Interest Statement

The authors have declared no competing interest.

