## Supplemental Figures for "Rat somatic genome editing enables ER+ breast cancer modeling"

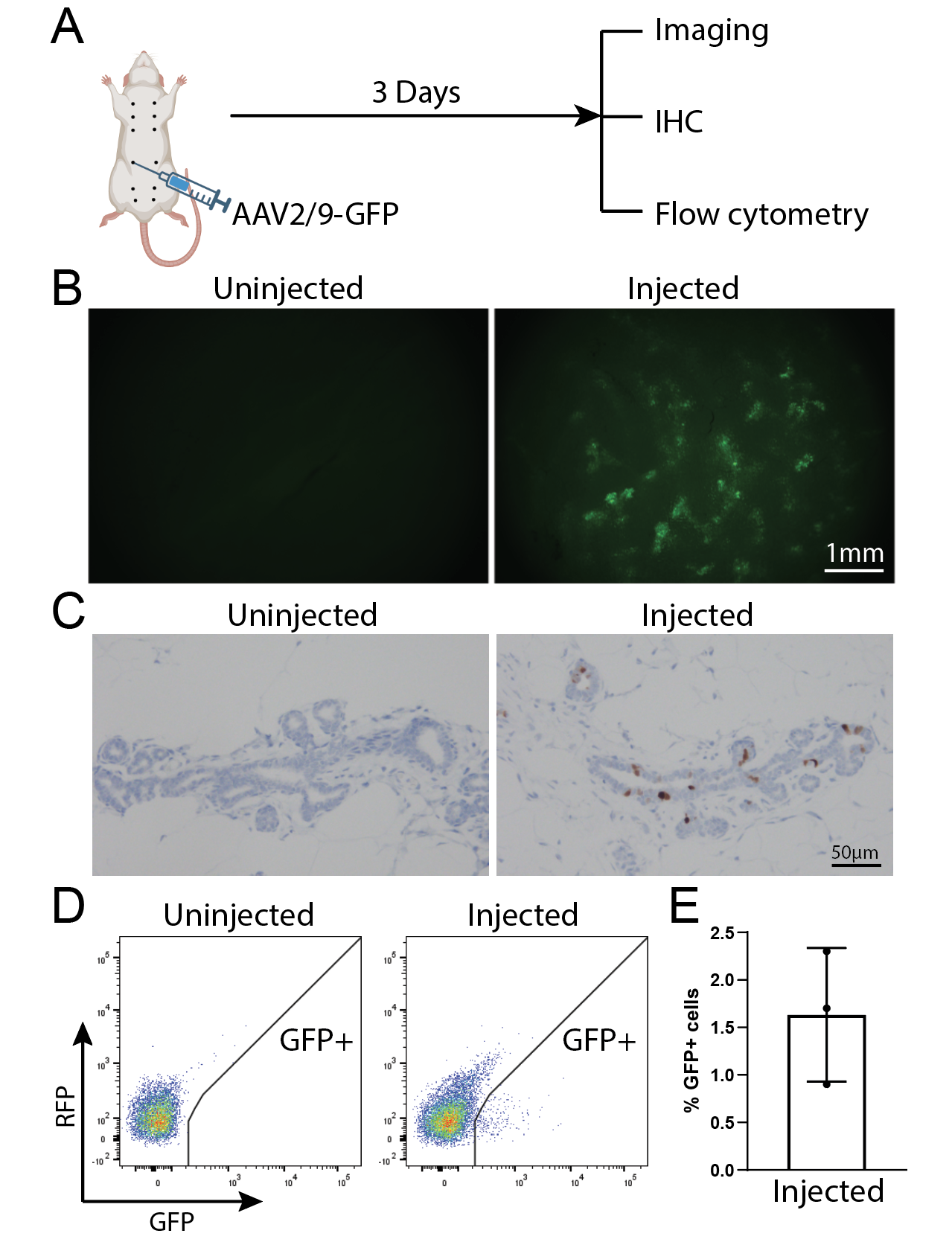

**Figure S1. Confirmation of cellular tropism of AAV2/9 in rat mammary epithelial cells.**

(A) Schematic representation of experimental design.

(B) Representative fluorescent microscope images of an AAV2/9-GFP-infected rat mammary glands compared to a non-infected control mammary gland.

(C) Immunohistochemical staining of GFP of AAV2/9-GFP-infected rat mammary glands and a non-infected control mammary gland.

(D) Representative flow cytometry analysis of single-cell suspensions prepared from an AAV2/9-GFP-infected rat mammary gland and a non-infected control mammary gland.

(E) Quantification of the percentage of infected cells. Cas9 rats, 8~9 weeks, n=3, virus dose: 3.75x10^9^gc/gland.

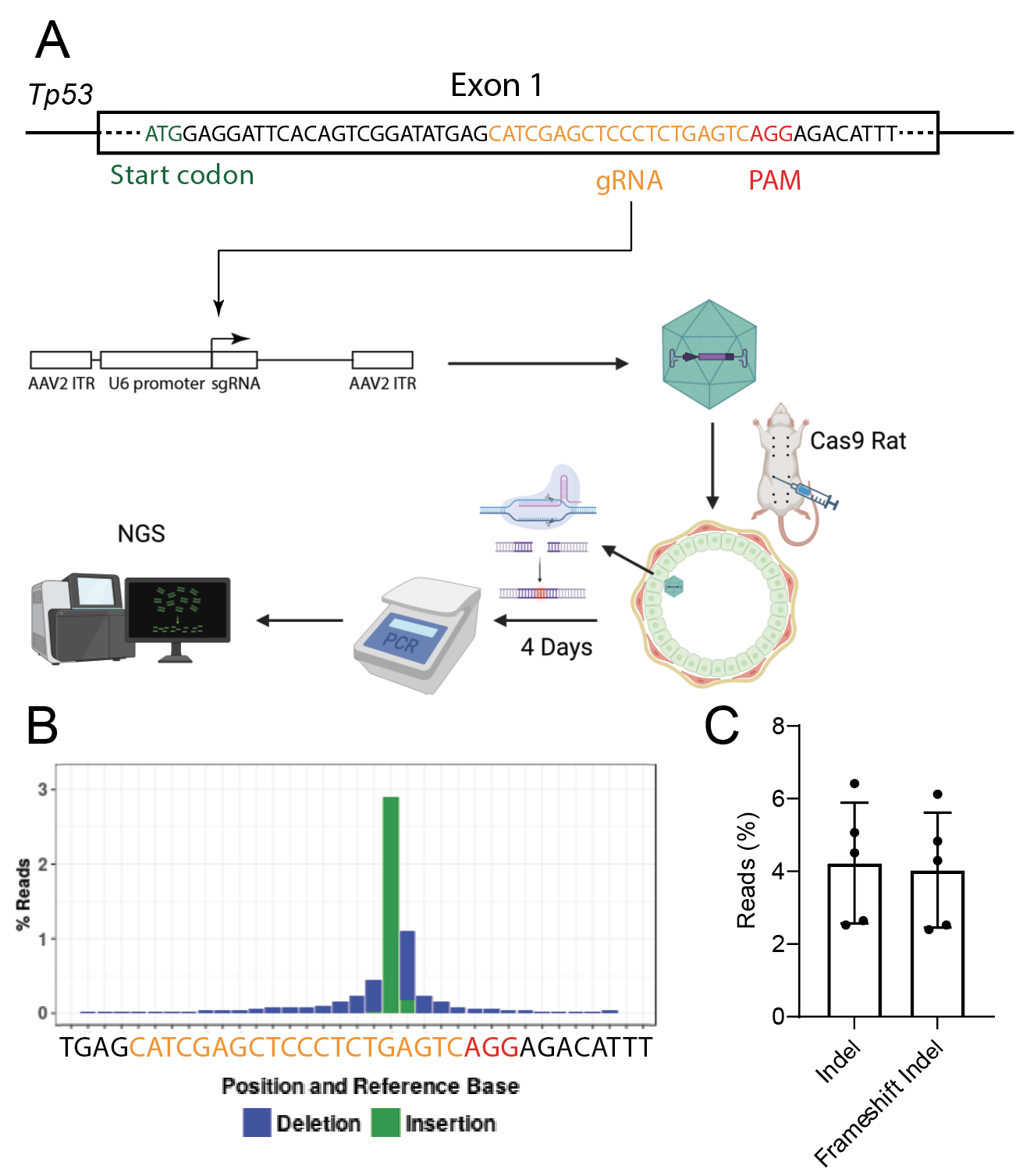

**Figure S2.** **Confirming somatic editing in rat mammary glands.**

(A) Schematic representation of experimental design to test somatic editing in rat mammary glands.

(B) Indel frequencies at the edited region, detected by amplicon sequencing of the targeted genomic region.

(C) Percentages of total indel alleles and frameshift indel mutations resulting from editing.

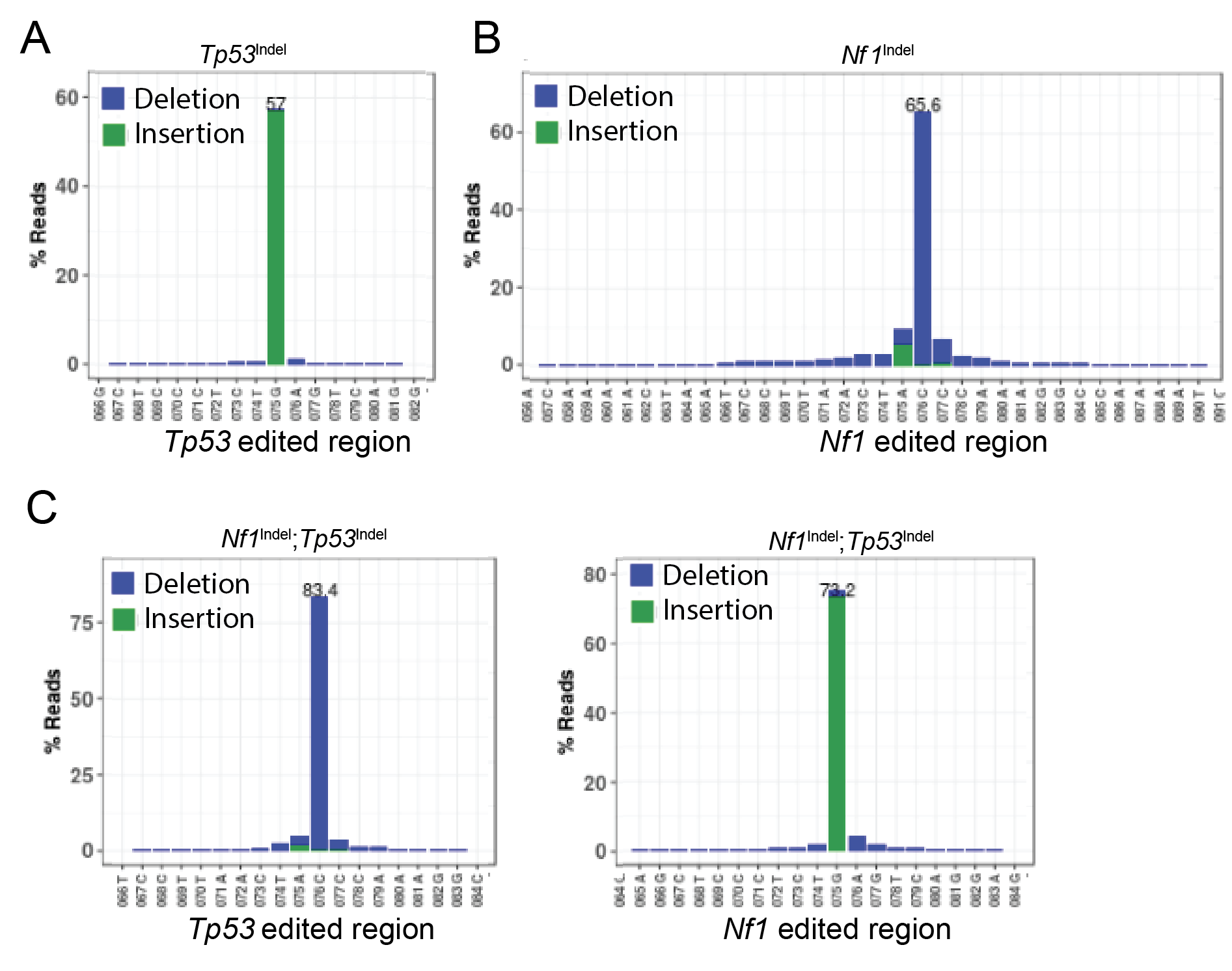

**Figure S3. Indel profiles of edited genome regions in mammary tumors induced by somatic indel editing in rats.**

(A) Representative indel profile of edited *Tp53* genome region in a mammary tumor induced by somatic editing of *Tp53* in a rat.

(B) Representative indel profile of edited *Nf1* genome region in a mammary tumor induced by somatic editing of *Nf1* in a rat.

(C) Representative indel profile of the edited *Tp53* and *Nf1* genome regions in a mammary tumor induced by somatic editing of *Tp53* and *Nf1* in a rat.

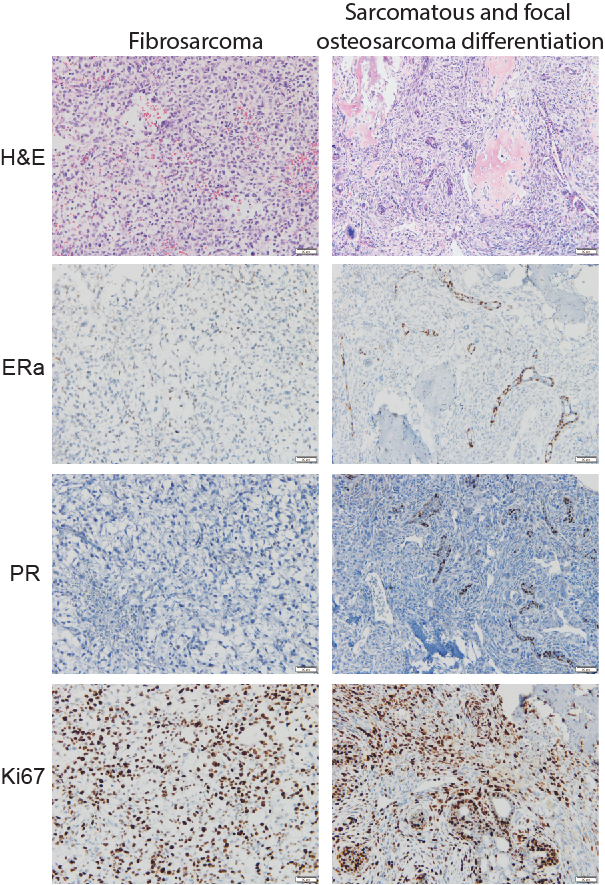

**Figure S4. Histopathological variants of *Tp53*^Indel^ tumors. Two of the five the *Tp53*^indel^ tumors examined are not typical ductal carcinomas.** One shows fibrosarcoma morphology (A). Another shows metaplastic differentiation of osteosarcoma (B).

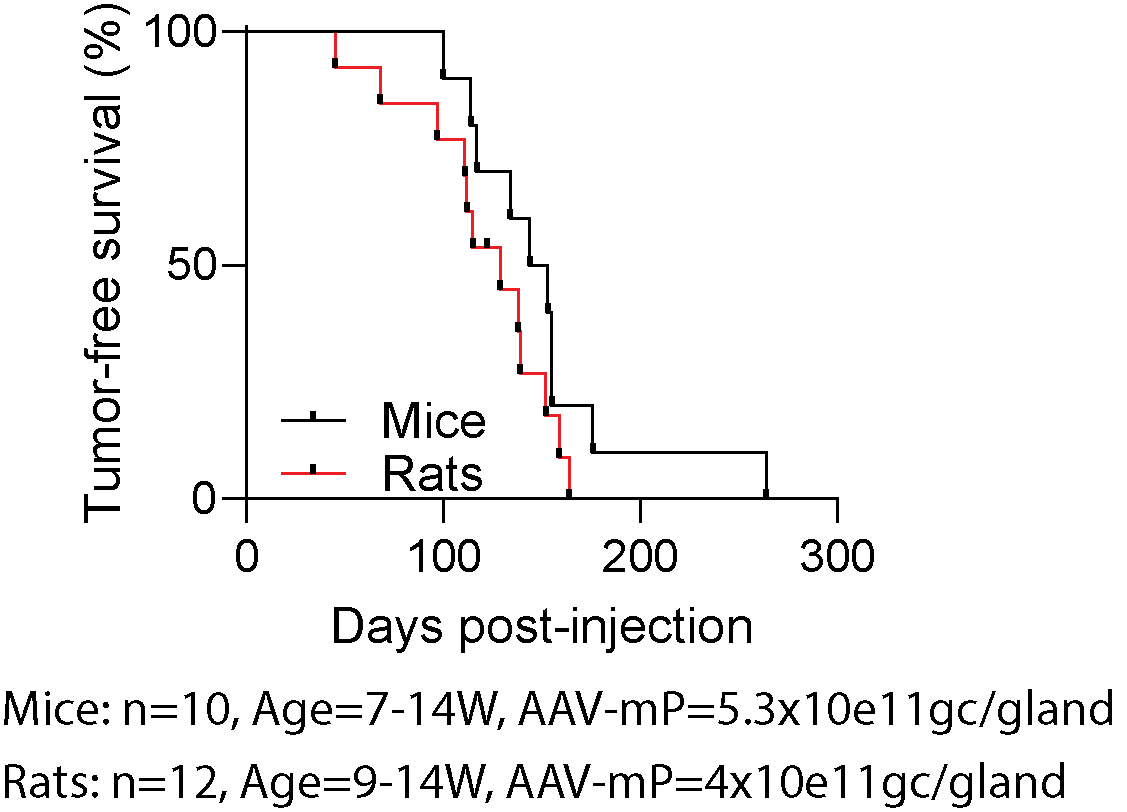

**Figure S5. *Pik3ca*^H1047R^ tumors in both mice and rats.** Kaplan-Meier tumor survival curves of mammary tumors induced by the same *Pik3ca*^H1047R^ somatic editing in mouse (black line) and rat (red line) mammary glands, respectively.

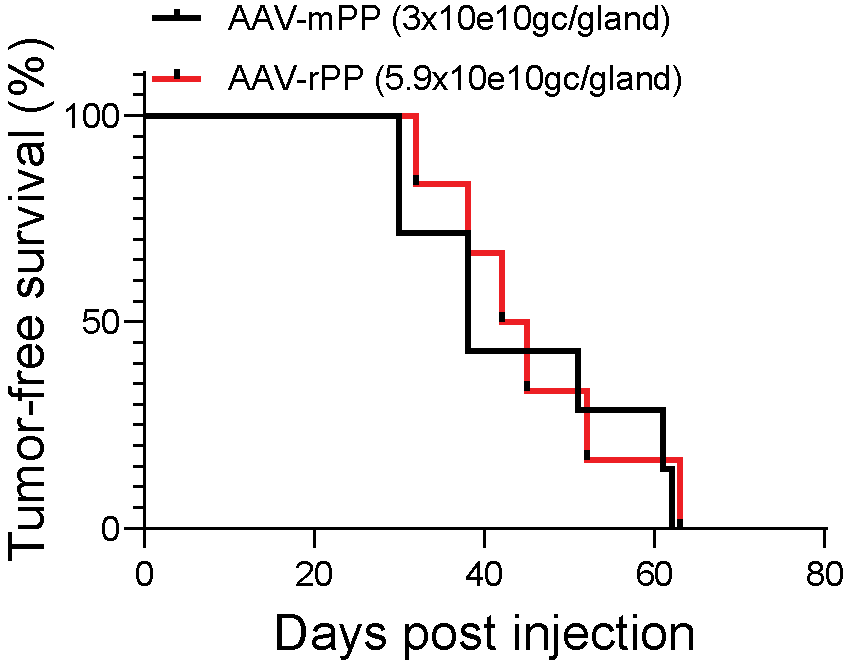

**Figure S6 *Tp53^Indel^*/*Pik3ca*^H1047R^ tumors in mice vs. rats.** Tumor-free survivals of mice (black line) and rats (red line) injected with the indicated AAV virus to introduce both *Tp53Indel* and *Pik3ca*^H1047R^.
